# Sparse dynamic graphical models for longitudinal non-Gaussian data

**DOI:** 10.64898/2026.09.25.754276

**Authors:** Yuhan Zheng, Claudia Solis-Lemus

**Affiliations:** Department of Statistics, University of Wisconsin-Madison; Wisconsin Institute for Discovery, University of Wisconsin-Madison

**Keywords:** Dynamic graphical model, Latent Gaussian model, Non-Gaussian longitudinal data, Monte Carlo EM, Microbiome data

## Abstract

Longitudinal non-Gaussian data are common in biomedical and microbiome studies, where responses may not be normally distributed. However, many existing dynamic graphical models are developed for Gaussian responses, while many non-Gaussian graphical models focus on cross-sectional dependence and do not explicitly estimate temporal transition effects. We propose a latent Gaussian dynamic graphical model for longitudinal non-Gaussian responses, including binary, count, and compositional outcomes. The proposed framework jointly estimates the contemporaneous conditional dependence network and the lag-one temporal dependence structure. We further develop a penalized Monte Carlo expectation maximization procedure for estimation. Simulation studies across multiple graph structures and outcome types show that the proposed method can recover both dependence structures, with more stable recovery for the lag-one temporal dependence structure and more structure-dependent recovery for the contemporaneous conditional dependence network. We apply the proposed method to longitudinal data from the Integrative Human Microbiome Project (HMP2) using a multinomial observation model. The results reveal persistent lag-one temporal dependence and contemporaneous conditional dependence among dominant microbial taxa, demonstrating the utility of the proposed framework for dynamic network analysis of longitudinal non-Gaussian biomedical data.

## 1. Introduction

Longitudinal data are common in biomedical studies, where repeated measurements are collected from the same subjects over time. In many applications, the responses are non-Gaussian: disease indicators may be binary, sequencing reads are counts, and microbiome profiles are often observed as compositional counts across microbial taxa (Lloyd-Price *and others*, 2019; Xia *and others*, 2013; Inouye *and others*, 2017). Such data can exhibit complex dependence structures, including contemporaneous conditional associations among variables and temporal dependence across adjacent time points (Abegaz and Wit, 2013).

Graphical models provide a useful framework for characterizing conditional dependence among multiple variables (Yuan and Lin, 2007; Friedman *and others*, 2008). A dynamic graphical model should distinguish contemporaneous conditional dependence from temporal dependence. The contemporaneous network describes conditional associations among variables measured at the same time point. The temporal transition structure characterizes how variables at one time point predict variables at the next time point. Separating these two components is important because two variables may be associated within the same time point without having a strong temporal relationship, and vice versa (Abegaz and Wit, 2013; Basu and Michailidis, 2015).

Many existing dynamic graphical models are formulated under Gaussian assumptions and are therefore not directly applicable to longitudinal binary, count, or compositional data (Abegaz and Wit, 2013; Epskamp *and others*, 2018). Graphical models for non-Gaussian or mixed data have also been developed using exponential family distributions and conditional modeling approaches. However, these methods primarily characterize contemporaneous dependence and do not explicitly model temporal transitions (Yang *and others*, 2015; Lee and Hastie, 2015). More recently, methods for multivariate count time series have incorporated both temporal dynamics and contemporaneous conditional dependence (Sathish *and others*, 2025; Kim *and others*, 2026). Nevertheless, these methods are specifically designed for count data and do not provide a unified framework for other common non-Gaussian outcomes, such as binary and multinomial observations. Consequently, there remains a need for a flexible framework that accommodates multiple types of longitudinal non-Gaussian responses while jointly estimating contemporaneous and temporal dependence structures.

In this paper, we propose a latent Gaussian dynamic graphical model for longitudinal non-Gaussian observations. This framework links a latent Gaussian state process to binary, count, or composition observation models. Within the latent process, the precision matrix characterizes contemporaneous conditional dependence, whereas the transition matrix captures lag-one temporal dependence; they jointly estimate these two dependence structures from longitudinal non-Gaussian observations. We evaluate its performance through simulation studies covering multiple graph structures and outcome types and demonstrate its practical utility through an application to longitudinal data from the Integrative Human Microbiome Project (HMP2) (Lloyd-Price *and others*, 2019) using a multinomial observation model.

## 2. Methods

### 2.1 Latent Gaussian dynamic graphical model

We consider a class of multivariate longitudinal outcome models built on a common latent Gaussian state-space structure (Durbin and Koopman, 2012). This construction is motivated by sparse time series chain graphical models for dynamic network estimation, in which temporal dependence and contemporaneous conditional dependence are represented by separate model parameters (Abegaz and Wit, 2013). Let *Y*_*it*_ denote the observed response for subject *i* = 1, …, *N* at time *t* = 1, …, *T* . The form of *Y*_*it*_ depends on the outcome type (binary, count, or composition) and is specified in Section 2.2. Across all outcome types, we introduce a *p*-dimensional latent Gaussian state vector *Z*_*it*_ ∈ ℝ^*p*^, which evolves according to a first-order vector autoregressive model (VAR(1)):

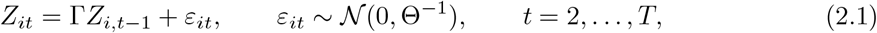

or equivalently,

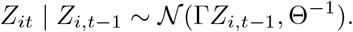

Here, Γ ∈ ℝ^*p*×*p*^ is the temporal transition matrix and Θ ∈ ℝ^*p*×*p*^ is the innovation precision matrix, with Θ ≻ 0. The matrix Γ encodes temporal dependence among latent features across adjacent time points: an entry Γ_*jk*_≠ 0 indicates that the conditional mean of the *j*-th latent feature at time *t* depends on the *k*-th latent feature at time *t*−1, after accounting for the remaining latent features at time *t* − 1. In contrast, the off-diagonal entries of Θ encode contemporaneous conditional dependence among the innovations at time *t*. Thus, Γ defines the temporal dependence network, whereas Θ defines the contemporaneous conditional dependence network. For the initial state, we assume the following:

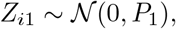

where *P*_1_ is chosen as a fixed diffuse positive definite covariance matrix to represent weak prior information about the initial latent state (Durbin and Koopman, 2012). Since the initial state density does not involve (Γ, Θ), it is omitted from the maximization step derivations. However, it is included when sampling the latent trajectories in the Monte Carlo Expectation step (E-step). (see Section 2.3).

This latent dynamic graphical model serves as the Gaussian distribution for all observation models considered in this work. Following latent Gaussian hierarchical constructions for non-Gaussian multivariate responses (Shen and Solís-Lemus, 2026), binary, count, and compositional outcomes are connected to the latent state through different observation layers. The observation model determines the form of the expectation approximation, while the latent transition model and the penalized maximum step for (Γ, Θ) are shared across outcome types. For binary and count outcomes, each of the *p* components of *Y*_*it*_ is linked to the corresponding component of *Z*_*it*_. For composition outcomes with *C* categories, *Y*_*it*_ is a single *C*-category count vector, and we use a baseline-category representation with *p* = *C* − 1.

### 2.2 Observation models

We model non-Gaussian outcomes through observation models conditional on a latent Gaussian state. This strategy follows the hierarchical extensions, where a Gaussian dependence structure is combined with appropriate observation layers for binary, count, and composition responses (Shen and Solís-Lemus, 2026). Specifically, we use a probit latent-variable representation for binary outcomes (Albert and Chib, 1993; Shen and Solís-Lemus, 2026), a lognormal-Poisson construction for count outcomes (Aitchison and Ho, 1989; Shen and Solís-Lemus, 2026), and a normal-logistic multinomial model for compositional count outcomes (Xia *and others*, 2013; Shen and Solís-Lemus, 2026). These constructions preserve the Gaussian latent state-space structure while allowing different observation types.

#### 2.2.1 Binary outcomes

*Probit observation model*. For binary responses, we use a probit observation model with Gaussian data augmentation (Albert and Chib, 1993; Shen and Solís-Lemus, 2026). We observe a vector of *p* binary responses

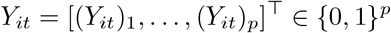

for subject *i* = 1, …, *N* at time *t* = 1, …, *T* . Conditional on the latent state *Z*_*it*_, the components of *Y*_*it*_ are assumed independent, with

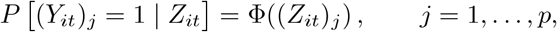

where Φ(·) denotes the standard normal cumulative distribution function. Hence, the conditional likelihood is

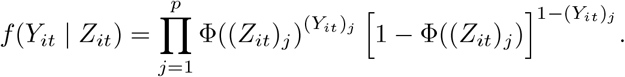

Although the binary components are conditionally independent given *Z*_*it*_, dependence among the observed responses is induced through the latent Gaussian state process.

##### Data augmentation and Gaussian state-space representation

Estimation (Section 2.3) requires sampling the latent states given the observed outcomes, which is not directly feasible under the probit model. To make this tractable, we introduce an auxiliary Gaussian vector 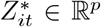 following the data augmentation strategy of Albert and Chib (1993); Holmes and Held (2006). We distinguish between the latent state *Z*_*it*_, which follows the VAR(1) transition model, and the auxiliary variable 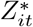, which is introduced only for data augmentation in the observation equation. Specifically,

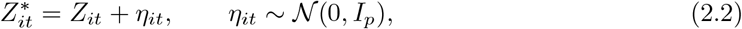

and

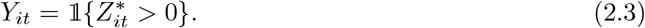

Integrating out 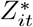 recovers the probit model *P{*(*Y*_*it*_)_*j*_ = 1 | *Z*_*it*_*}* = Φ ((*Z*_*it*_)_*j*_), and the unit variance of *η*_*it*_ fixes the scale of the latent state. By equations (2.2) and (2.3), the components of 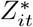 are conditionally independent truncated normal variables given *Z*_*it*_ and *Y*_*it*_:

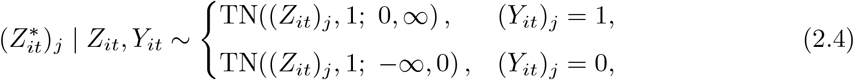

where TN(*µ, σ*^2^; *a, b*) denotes a normal distribution with mean *µ* and variance *σ*^2^, truncated to (*a, b*).

Conditional on the augmented observations 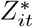, the binary model becomes a linear Gaussian state-space model with VAR(1) (2.1) and observation equation (2.2). In this augmented model, the observation matrix is the identity matrix and the observation noise covariance is *I*_*p*_. This augmentation converts the binary observation model into a conditionally Gaussian state space model, enabling latent state sampling via standard Gaussian simulation smoothing methods (Carter and Kohn, 1994; Durbin and Koopman, 2002).

#### 2.2.2 Count outcomes

##### Lognormal-Poisson observation model

We use a lognormal-Poisson observation model, a standard latent Gaussian construction for multivariate count data (Aitchison and Ho, 1989; Shen and Solís-Lemus, 2026). We observe a vector of *p* nonnegative integer counts

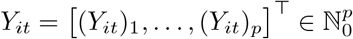

for subject *i* = 1, …, *N* at time *t* = 1, …, *T* . Conditional on *Z*_*it*_, the components of *Y*_*it*_ are assumed independent, with

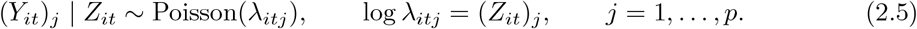

Hence, the conditional likelihood is

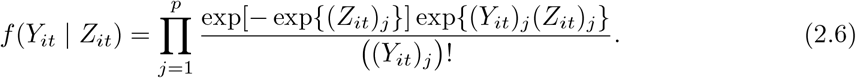

Although the count components are conditionally independent given *Z*_*it*_, dependence among the observed counts is induced through the latent Gaussian state process. In addition, the latent Gaussian layer allows the marginal count distribution to accommodate overdispersion relative to a standard Poisson model (Inouye *and others*, 2017). When counts are observed with varying exposure, such as library size in sequencing data, a known offset can be added to log *λ*_*itj*_ in equation (2.5) without changing the estimation procedure.

##### Gaussian approximation and importance sampling

Under the count observation model in equation (2.5), the smoothing distribution of the latent states conditional on the observed counts is non-Gaussian, and the moments required in the E-step are not available in closed form. We therefore construct a linear Gaussian approximation to the count state space model using the iterative procedure implemented in `KFAS` (Helske, 2017). Let 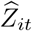 denote the conditional mode under the current parameter estimates and define 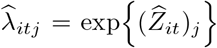 . The working represents the count observation likelihood Gaussian observation

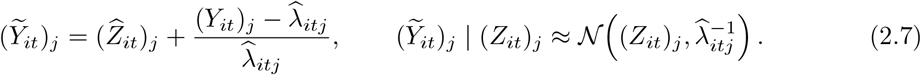

Combining equation (2.7) with the latent VAR(1) transition model gives an approximate Gaussian state space model, from which latent state trajectories are generated using Gaussian simulation smoothing (Durbin and Koopman, 2002).

Because these trajectories are generated under the Gaussian approximation, an importance correction is applied using the likelihood ratio

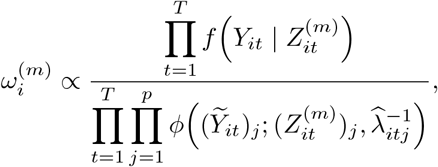

where 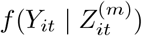 is the exact Poisson likelihood in equation (2.6), and *ϕ*( · ; *µ, σ*^2^) denotes the Gaussian density with mean *µ* and variance *σ*^2^. These likelihood ratios are normalized across the sampled trajectories and used to estimate the conditional moments involving 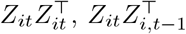, and 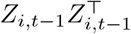 required for updating Γ and Θ. Thus, the Gaussian approximation enables efficient latent state simulation, while the importance correction targets the original count observation model.

#### 2.2.3 Compositional outcomes

##### Normal-logistic multinomial observation model

For compositional outcomes, let *Y*_*it*_ = ((*Y*_*it*_)_1_, …, (*Y*_*it*_)_*C*_)^*T*^ denote the *C*-category multinomial response for subject *i* = 1, …, *N* at time *t* = 1, …, *T*, where each (*Y*_*it*_)_*c*_ ⩾ 0 and 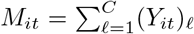 is the total count at time *t*. We use the *C*-th category as the baseline and represent the latent logit vector by

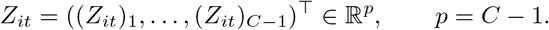

The baseline category fixes the location of the latent vector, since equation (2.8) is invariant to adding a common constant to all components of *Z*_*it*_. Following the normal logistic multinomial construction for compositional count data (Liu, 2001; Shen and Solís-Lemus, 2026), the category probabilities are defined through the baseline category compositional logit link:

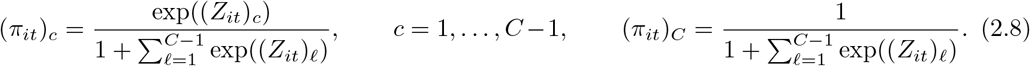

Equivalently,

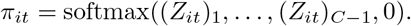

Conditional on the latent state *Z*_*it*_, the observed counts follow

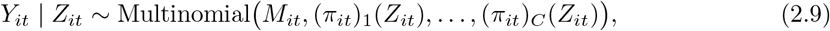

with conditional probability mass function

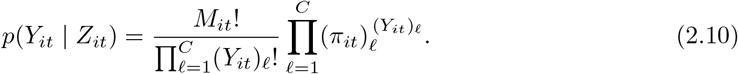

##### Laplace approximation and pseudo-Gaussian state-space representation

Under the normal-logistic compositional observation model, the conditional distribution of *Z*_*it*_ given *Y*_*it*_ is not Gaussian, and its moments are not available in closed form. We therefore construct a Laplace pseudo-Gaussian state-space approximation for the Monte Carlo E-step (Tierney and Kadane, 1986). Up to an additive constant, the conditional log likelihood contribution of *Y*_*it*_ given *Z*_*it*_ is

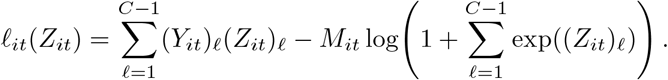

Let 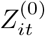 be a current reference point, and let 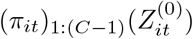 denote the first *C* − 1 category probabilities evaluated at 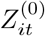. The score vector and negative Hessian matrix of *l*_*it*_(*Z*_*it*_) at 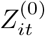 are

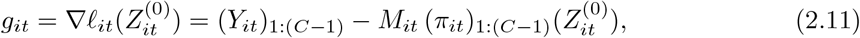

and

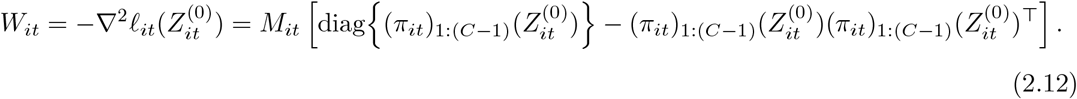

A second-order Taylor expansion around 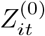 gives

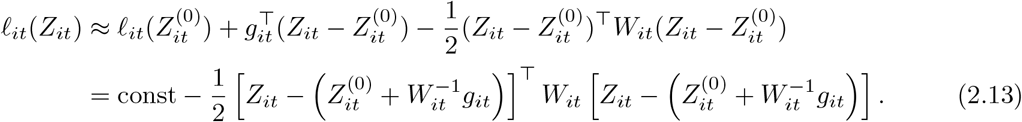

Therefore, defining the pseudo-observation

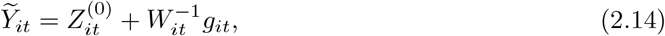

We obtain the pseudo-Gaussian approximation

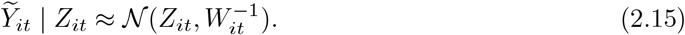

Combining equation (2.15) with the latent VAR(1) transition model (equation (2.1)) yields an approximate Gaussian state-space model. This Gaussian approximation provides a tractable proposal distribution for latent-state sampling using Kalman filtering and Gaussian simulation smoothing (Carter and Kohn, 1994; Durbin and Koopman, 2002).

To correct for the approximation and target the exact multinomial observation model, we use importance reweighting of the sampled latent trajectories (Liu, 2001). For a sampled trajectory 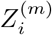 drawn from the pseudo-Gaussian proposal distribution, the unnormalized importance weight is proportional to

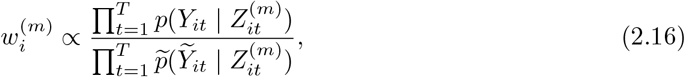

where 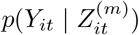 is the exact multinomial likelihood in (2.10), and 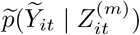 is the pseudo-Gaussian likelihood in equation (2.15). The normalized weights are then used to approximate the conditional expectations of the complete data sufficient statistics.

### 2.3 Estimation via penalized Monte Carlo EM

Direct maximization of the observed-data likelihood is difficult because the latent states *{Z*_*it*_*}* are unobserved, and the observation models are non-Gaussian. We therefore estimate (Γ, Θ) using a penalized Monte Carlo EM framework (Wei and Tanner, 1990; Green, 1990). Let *l*_*c*_(Γ, Θ) denote the complete data log likelihood, where the complete data include the observed outcomes, the latent states, and any outcome-specific augmentation variables introduced in the Monte Carlo E-step when applicable. At iteration *k*, the *Q*-function is

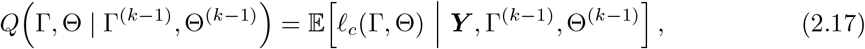

where

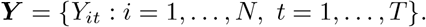

In practice, this conditional expectation is approximated using Monte Carlo samples from the conditional distribution of the latent trajectories given the observed outcomes.

To induce sparsity in both the temporal transition matrix and the innovation precision matrix, the maximization step (M-step) maximizes the penalized criterion

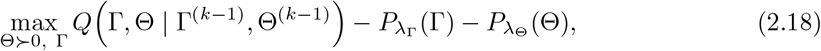

where 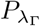 (Γ) and 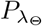 (Θ) are sparsity inducing penalty functions with tuning parameters *λ*_Γ_ and *λ*_Θ_, respectively (Fan and Li, 2001; Abegaz and Wit, 2013; Basu and Michailidis, 2015). Because the initial-state distribution is fixed and does not involve (Γ, Θ), its contribution can be ignored in the M-step. Up to additive constants that do not depend on (Γ, Θ), the complete data log likelihood can be decomposed as

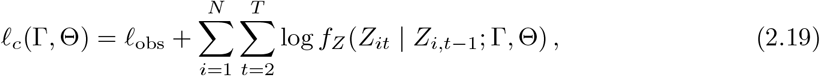

where *l*_obs_ collects the contribution from the observation model and any outcome-specific augmentation variables, and *f*_*Z*_(· | ·; Γ, Θ) is the Gaussian latent transition density induced by equation (2.1). The observation contribution determines the conditional distribution of the latent states given the observed outcomes in the E-step. However, conditional on the latent states and augmentation variables, *l*_obs_ does not contain (Γ, Θ). Therefore, it does not affect the algebraic form of the M-step objective for these parameters.

For notational convenience, we define

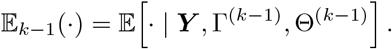

Then the transition part of the *Q*-function is given by

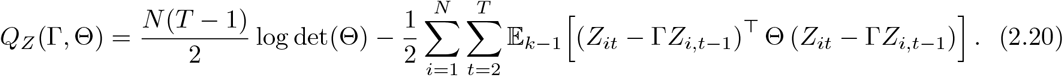

Using the trace identity *x*^⊤^*Ax* = tr(*Axx*^⊤^), the quadratic term can be written as

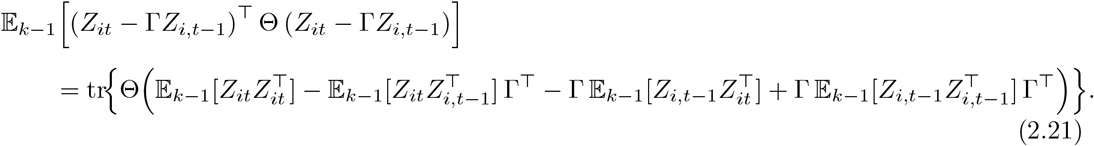

Hence, the M-step depends on the conditional second moments

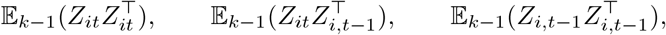

which are approximated in the outcome-specific Monte Carlo E-step. For notational convenience, we define the aggregated sufficient statistics

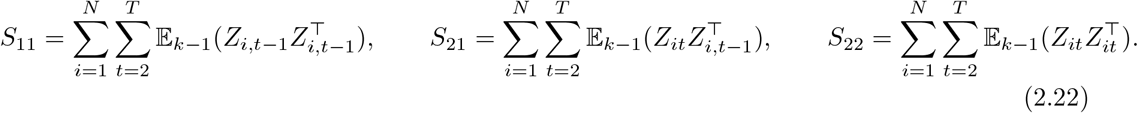

These quantities are computed in the E-step and passed to the penalized M-step. In terms of *S*_11_, *S*_21_, and *S*_22_, the transition part of the *Q*-function in (2.20) can be written as

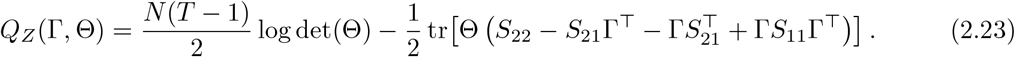

#### Monte Carlo E-step for each outcome type

The Monte Carlo E-step is outcome-specific because the conditional distribution of the latent trajectories given the observed outcomes depends on the observation model. However, once samples from the latent trajectories are obtained, all outcome types lead to the same complete data sufficient statistics *S*_11_, *S*_21_, and *S*_22_ defined in equation (2.22). Thus, the subsequent penalized M-step (next subsection) is shared across binary, count, and compositional outcomes.

For binary outcomes, the probit data augmentation of Section 2.2.1 introduces auxiliary Gaussian variables 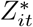 (Holmes and Held, 2006; Shen and Solís-Lemus, 2026). Conditional on these augmented observations, the model becomes a linear Gaussian state-space model, so latent trajectories can be sampled using Gaussian simulation smoothing (Carter and Kohn, 1994; Durbin and Koopman, 2002). For count outcomes, the lognormal-Poisson observation model yields a non-Gaussian conditional distribution of the latent states given the observed counts (Aitchison and Ho, 1989; Shen and Solís-Lemus, 2026), which we sample using a linear Gaussian approximation to construct an importance proposal for estimating the conditional expectations required in the E-step. Since the real data application in this paper uses composition outcomes, we present the compositional E-step in detail below. The corresponding E-step algorithms for binary and count outcomes are provided in the Supplementary Material.

For composition outcomes, the E-step is implemented using a Laplace Gaussian proposal (Tierney and Kadane, 1986) and importance reweighting (Liu, 2001). For each subject *i* = 1, …, *N*, we first construct a pseudo-Gaussian state space approximation to the composition observation model, as in equation (2.15). The reference point 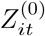 is chosen by an inner iteration: starting from an initial trajectory, we compute the pseudo observations 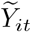 and precision matrices *W*_*it*_ at the current 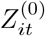, obtain the smoothed mean 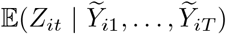 under the resulting Gaussian state space model, and set this mean as the new 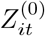, repeating until convergence. At convergence, 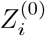 is the mode of the conditional distribution of the latent trajectory given *Y*_*i*_ as required by the Laplace approximation. After convergence of the inner loop, we draw latent trajectories from the conditional distribution of *Z*_*i*_ given the pseudo-observations under the Gaussian state-space model, using Gaussian simulation smoothing (Carter and Kohn, 1994; Durbin and Koopman, 2002). Importance weights are computed using the exact multinomial likelihood relative to the Gaussian pseudo-likelihood. Since the target conditional distribution and the proposal distribution share the same Gaussian transition density under the current parameter values, the transition contribution cancels in the importance ratio. The weighted samples are then used to estimate *S*_11_, *S*_21_, and *S*_22_. Implementation and numerical stabilization details for the multinomial E-step, including regularization of the pseudo-information matrices and stabilization of importance weights, are provided in the Supplementary Material. Algorithm 1 summarizes these steps. For notational convenience, define the row-stacked latent state matrices

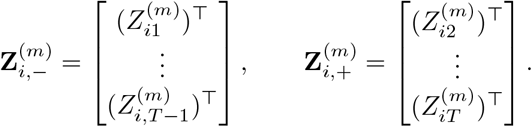

#### Penalized M-step shared across outcome types

Given the outcome-specific E-step estimations of *S*_11_, *S*_21_, and *S*_22_ in equation (2.22), we update (Γ, Θ) using a two-stage generalized M-step (Dempster *and others*, 1977). The first stage updates the transition matrix Γ through a penalized least squares working objective based on the latent VAR(1) transition model. The second stage constructs a residual covariance matrix using the updated Γ, and then updates the innovation precision matrix Θ by sparse precision estimation. Convergence is monitored using the normalized Frobenius norm in Γ and Θ, and the convergence diagnostic plots are provided in the Supplementary Materials. In the implementation, the Γ-update is solved using coordinate gradient descent with the smoothly clipped absolute deviation (SCAD) penalty (Gorstein *and others*, 2025), and the Θ-update is solved using the graphical lasso. Additional implementation details, including numerical stabilization, warm starts, and software routines, are provided in the Supplementary Material. Algorithm 2 summarizes the above steps.

##### Algorithm 1: Monte Carlo E-step for compositional outcomes

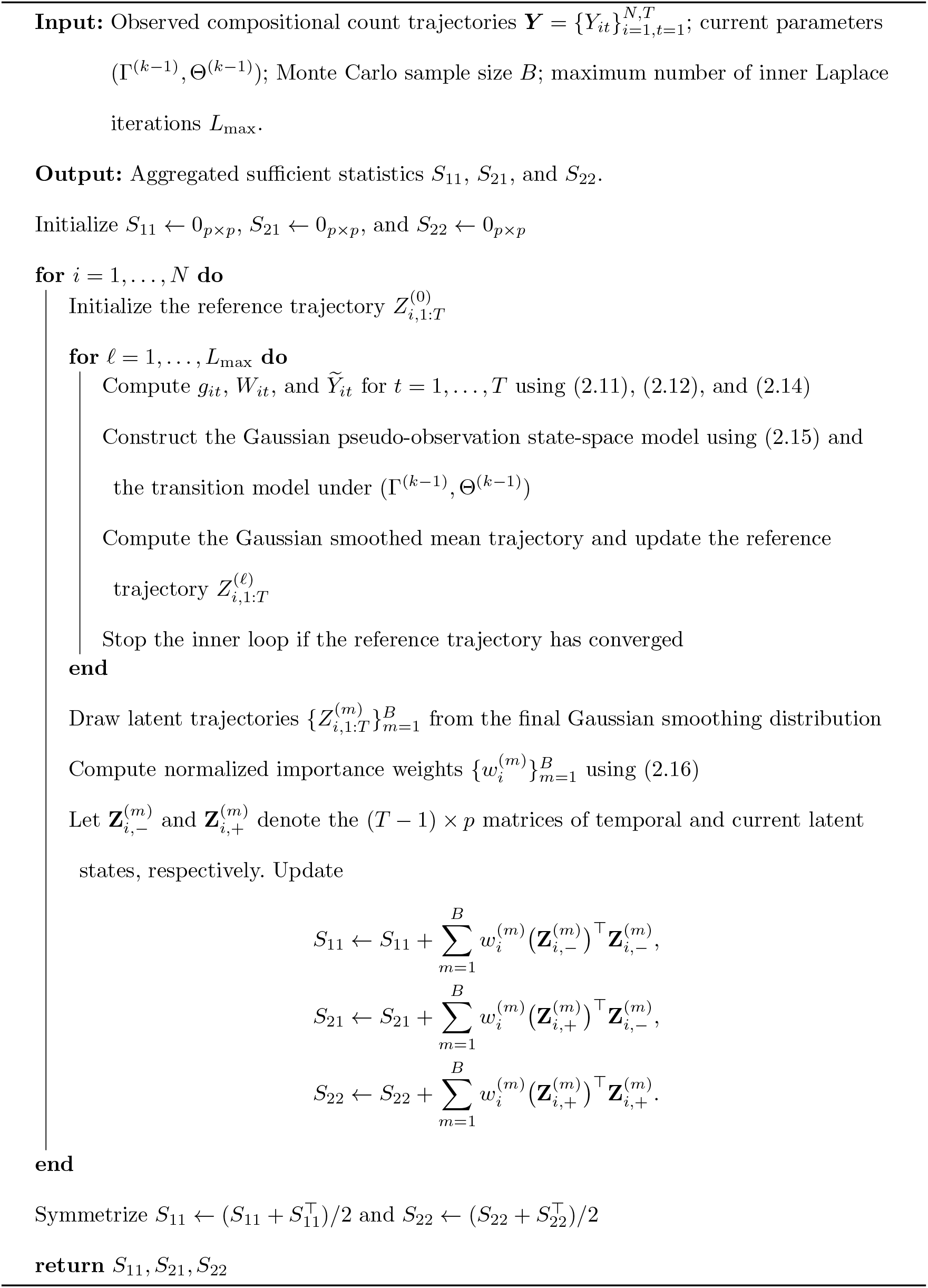

##### Algorithm 2: Penalized M-step and convergence check

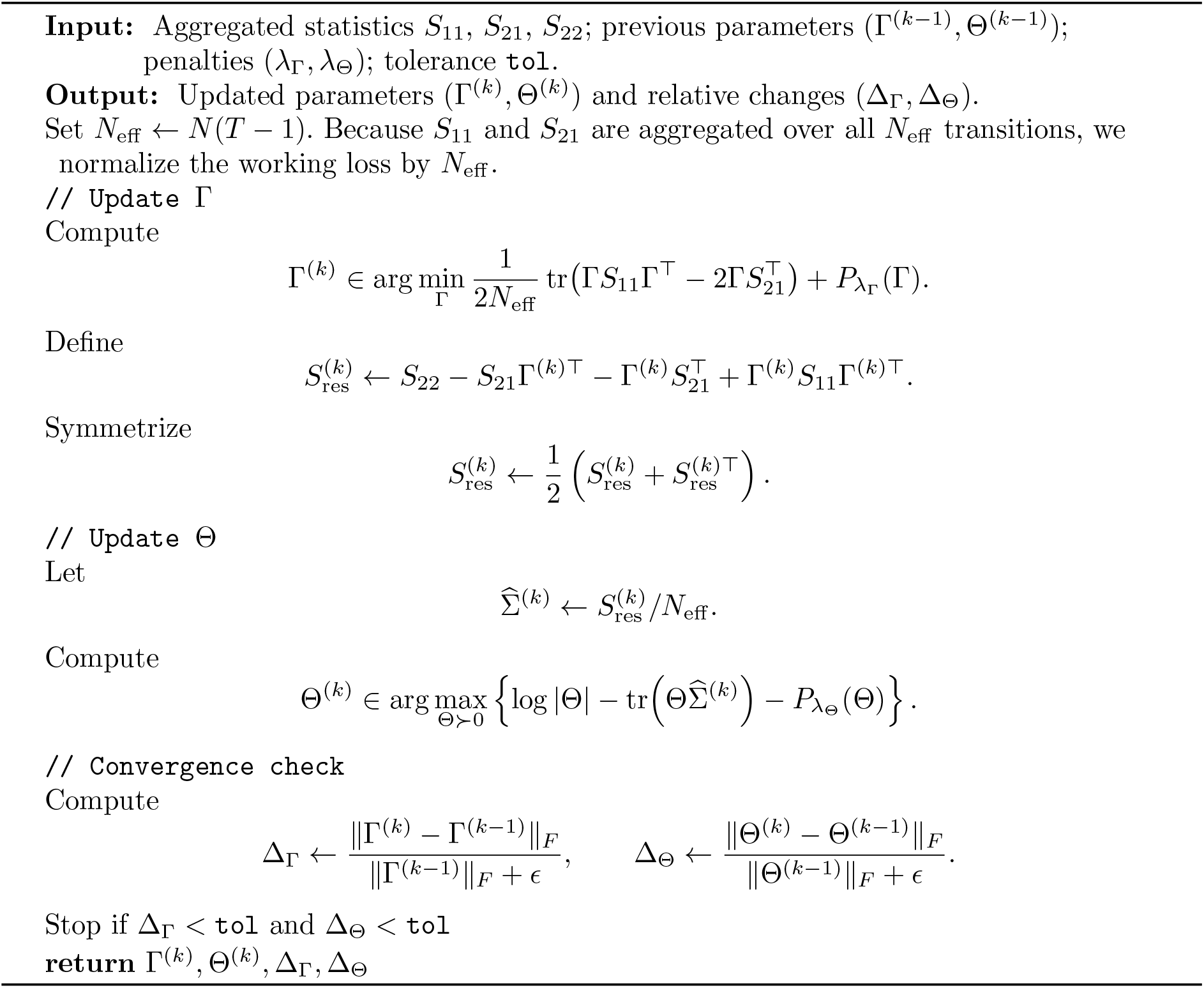

### 2.4 Model selection

The proposed estimation procedure involves two tuning parameters: *λ*_Γ_, which controls sparsity in the temporal transition matrix Γ, and *λ*_Θ_, which controls sparsity in the innovation precision matrix Θ. We use an EBIC criterion as the primary model selection criterion over a two-dimensional tuning grid. The EBIC was proposed as an extension of the classical BIC for model selection over large model spaces (Chen and Chen, 2008), and has been adapted to Gaussian graphical models for sparse precision matrix estimation (Foygel and Drton, 2010).

For each candidate tuning parameter pair (*λ*_Γ_, *λ*_Θ_), let 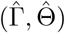 denote the final estimates, and let 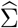 be the residual covariance matrix computed in Algorithm 2. We compute

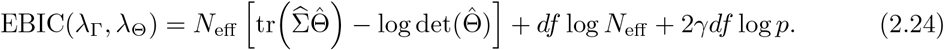

Here, *N*_eff_ is the effective sample size, *p* is the latent dimension, and *df* denotes the number of selected free parameters, computed as the number of nonzero entries in 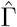 plus the number of nonzero off-diagonal entries in the upper triangular part of 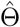. The parameter *γ* ∈ [0, 1] controls the strength of the additional high-dimensional complexity penalty; we set *γ* = 0.5, following the common choice used in EBIC-type graphical model selection (Foygel and Drton, 2010).

In practical applications, where the true network structure is unknown, the tuning parameters are selected by minimizing equation (2.24) over a candidate grid *G*:

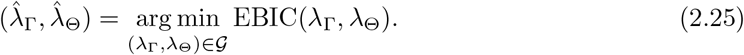

In the simulation study, where the true sparsity patterns are known, EBIC is used as the pilot selection criterion before the tuning parameter selection described in Section 3.2.

## 3. Simulation study

We considered six contemporaneous dependence structures, following the graph structures used in (Shen and Solís-Lemus, 2026). Some structures were specified through the covariance matrixΣ = Θ^−1^, whereas others were specified directly through the precision matrix Θ. When a constructed covariance or precision matrix was not positive definite, we added a small diagonal adjustment before inversion or simulation. Figure 1 and Table 1 illustrate the six contemporaneous network structures. Note that for the block structure, the variables were divided into two equally sized groups. Within each block, the covariance was set to 0.5, while the covariance between variables in different blocks was set to zero.

**Table 1:**
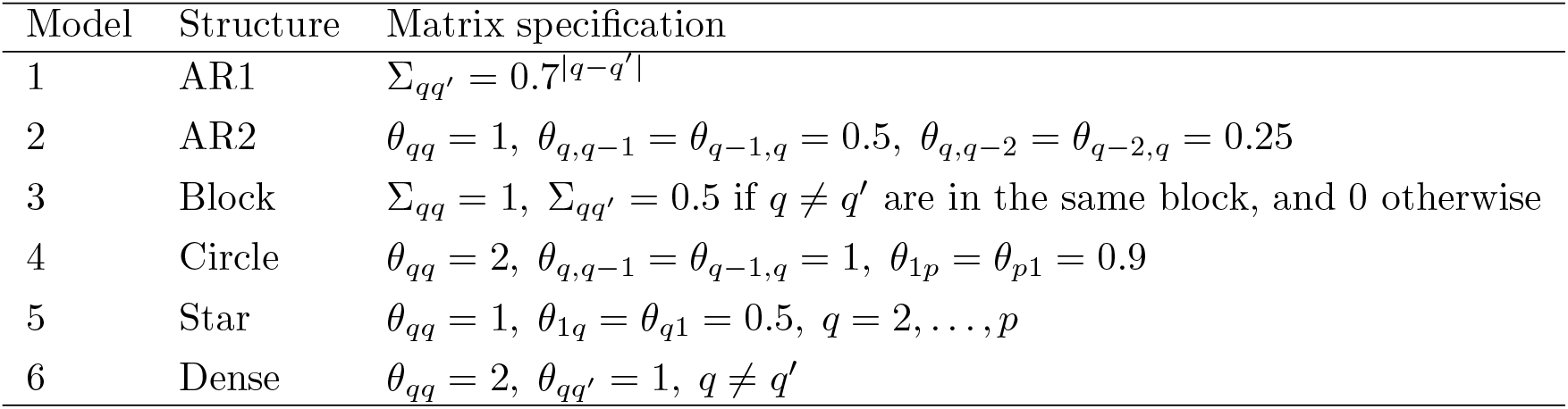
Six contemporaneous dependence structures used in the simulation study.

**Fig. 1:**
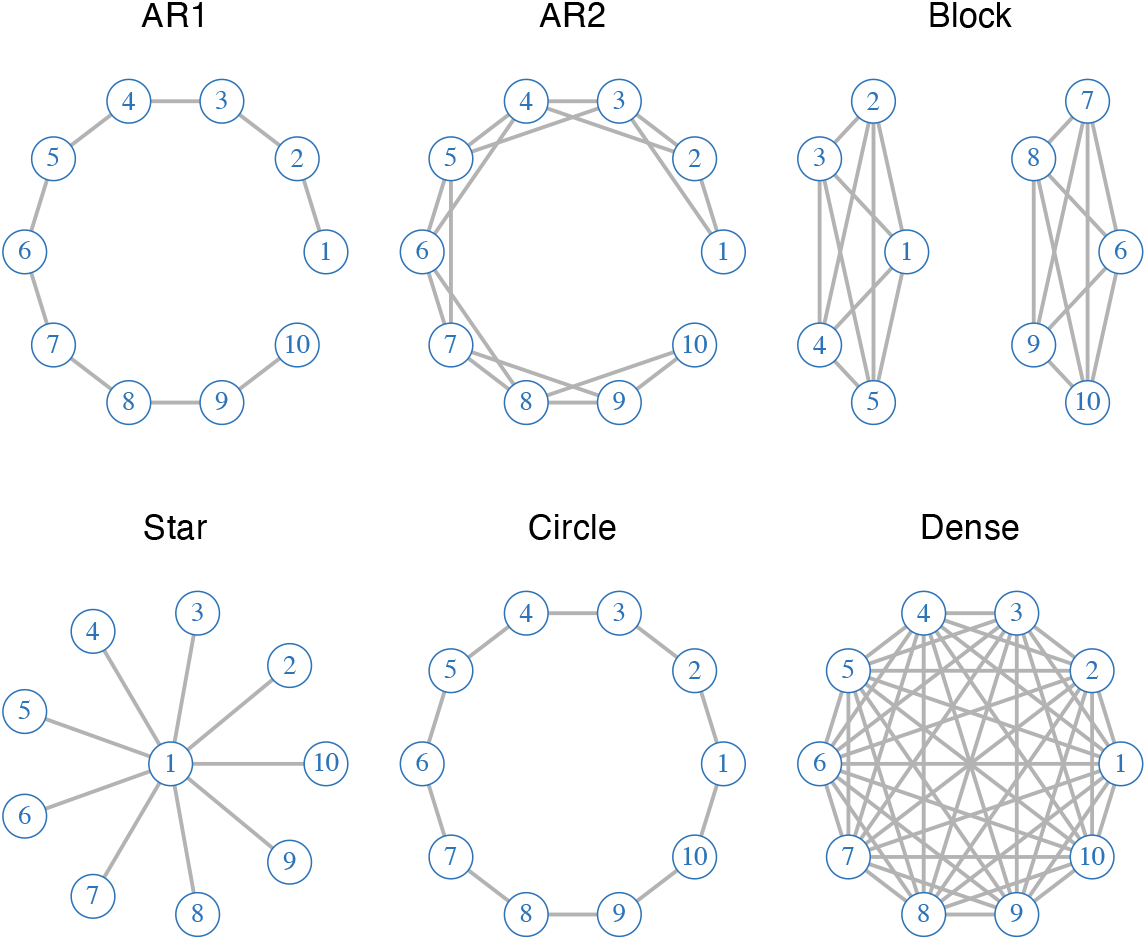
Six contemporaneous network structures used in the simulation study when *p* = 10. Table 1 shows the mathematical formulations of the covariance structures.

For each simulation replicate, we generated a sparse temporal transition matrix Γ ∈ ℝ^*p*×*p*^. Specifically, 20% of the entries were selected uniformly at random to be nonzero, and the nonzero entries were independently generated from a standard normal distribution. To ensure stability of the latent VAR(1) process, the resulting matrix was rescaled so that its spectral radius was strictly smaller than one.

Given the latent process *{Z*_*it*_*}*, we generated binary, count, and compositional outcomes according to the observation models described in Section 2.2. For the binary setting, we used the probit observation model

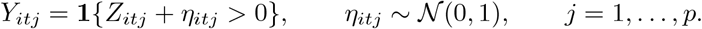

For the count setting, we generated conditionally independent counts with

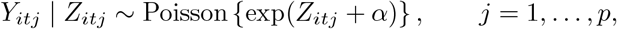

where *α* controls the marginal count level. For the compositional setting, we used *C* categories, with the *C*th category treated as the baseline. The category probabilities were generated by applying the softmax transformation to

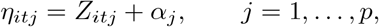

and *Y*_*it*_ | *Z*_*it*_ was then sampled from a multinomial distribution with total count *M*_*it*_.

### 3.1 Performance metrics

We evaluated support recovery for both the temporal transition matrix Γ and the contemporaneous precision matrix Θ. A nonzero entry was treated as a recovered edge. For Γ, all *p*^2^ entries were evaluated, including the diagonal entries which represent temporal self-dependence. For the symmetric precision matrix Θ, only the off-diagonal entries in the upper triangular part were evaluated. For each fitted model, an estimated entry was classified as nonzero if its absolute value exceeded a specified threshold 0.01. The estimated support was then compared with the true support to compute the numbers of true positives TP, true negatives TN, false positives FP, and false negatives FN. We summarized support recovery using sensitivity (SEN), specificity (SPE), and the Matthews correlation coefficient (MCC), and the *F*_1_ score:

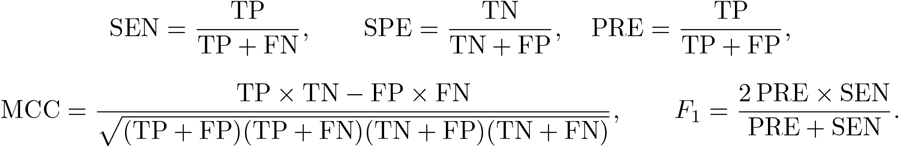

Higher values of SEN, SPE, and *F*_1_ indicate better support recovery, while MCC takes values in [−1, 1], with larger values indicating better agreement between the estimated and true sparsity patterns (Matthews, 1975; Powers, 2011).

### 3.2 Experimental settings and tuning parameter selection

We considered combinations of sample size, time length, and latent dimension given by

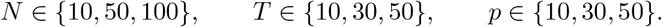

For each combination of (*N, T, p*), each outcome type, and each of the six contemporaneous dependence structures described above, we generated 100 independent simulation replicates. In all experiments, the Monte Carlo EM algorithm was run with a Monte Carlo sample size *B* = 100 at each iteration.

For each simulation setting, defined by a particular combination of (*N, T, p*), outcome type, and contemporaneous dependence structure, we conducted a separate pilot tuning procedure. The pilot dataset was generated under the same configuration but was not included among the 100 simulation replicates used to summarize support-recovery performance. During the pilot stage, the model was fitted over the two dimensional grid of regularization parameters

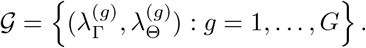

For each candidate pair, we computed the EBIC criterion described in Section 2.4. Because the true graph structures were known in the simulation study, we also computed MCC_Γ_ and MCC_Θ_ by comparing the estimated and true supports, using the same 0.01 threshold described in Section 3.1. We defined

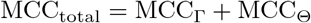

when both component specific MCC values were defined. For the Dense contemporaneous structure, all true off-diagonal entries of Θ were nonzero, so MCC_Θ_ was undefined. In this case, the available component score MCC_total_ = MCC_Γ_ was used during pilot selection.

The pilot selection proceeded in two stages. In the first stage, all candidate pairs were ranked by EBIC in ascending order, and the 10 pairs with the smallest EBIC values were retained. If two candidate pairs had identical EBIC values, the pair with the larger MCC_total_ was ranked first. In the subsequent selection stage, the retained candidate with the largest MCC_total_ was selected. If two or more retained pairs had the same MCC_total_, the pair with the smallest EBIC was selected. EBIC and MCC played complementary roles in this two-stage procedure. EBIC evaluates a candidate model through its likelihood while penalizing model complexity, thereby screening for models with a favorable balance between goodness of fit and parsimony. In contrast, MCC directly measures agreement between the estimated and true sparsity patterns in the simulation setting, but it does not incorporate likelihood fit or an explicit complexity penalty. Consequently, selection based only on EBIC need not identify the candidate with the highest support recovery accuracy, whereas selection based only on MCC would disregard model fit and complexity considerations. We therefore first used EBIC to restrict attention to a small set of competitive candidate models and then used MCC to distinguish their support recovery performance. This two-stage strategy balances likelihood model selection with recovery of the underlying graph structures under the simulation settings considered here.

For each simulation setting, the resulting tuning parameter pair 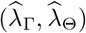 was held fixed and applied to all 100 independent simulation replicates. No additional tuning was performed within these evaluation replicates. The corresponding estimates 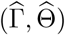 were then used to calculate the reported support recovery measures. Because the second stage relies on knowledge of the true graph structures, it was restricted to the simulation study. In the real data analysis, where the underlying network structures were unknown, the tuning parameter pair was selected by minimizing EBIC over the full candidate grid.

## 4. Simulation results

The simulation study addresses three main questions. First, we examine whether the proposed framework can recover the supports of the temporal dependence matrix Γ and the contemporaneous precision matrix Θ under different contemporaneous dependence structures. Second, we investigate how recovery performance changes with the sample size *N*, the number of time points *T*, and the latent dimension *p*. Third, we compare binary, count, and compositional outcomes to assess whether the observation model leads to systematic differences in support recovery performance.

### 4.1 Recovery under different graph structures

To assess support recovery under different structural patterns, we fix the baseline setting at (*N, T, p, B*) = (100, 10, 10, 100) and consider the six contemporaneous dependence structures (Figure 1). Note that for the Dense structure, every off-diagonal entry of the true precision matrix Θ is nonzero. This structure contains no true nonedges for Θ, so SPE, and MCC are undefined in this setting and are reported as not applicable. This degeneracy applies only to Θ; all recovery metrics remain well-defined for Γ, whose true support is sparse.

For each setting, results are averaged over 100 independent simulation replicates. Table 2 reports SEN, SPE, MCC, and *F*_1_ score for recovering the supports of Θ and Γ across the six structures and three outcome types at the baseline setting at (*N, T, p, B*) = (100, 10, 10, 100). Across the non-dense structures, recovery of Γ is more stable than recovery of Θ. Recovery of Θ depends more on the underlying contemporaneous structure. The bolded entries in Table 2 show that Block yields the numerically highest MCC_Θ_ for all three outcome types. This consistency suggests that the clustered dependence represented by the Block structure is favorable for contemporaneous support recovery under the baseline setting. The structure associated with the highest MCC_Γ_ is different from that associated with the highest MCC_Θ_. For the count outcome, AR2 yields the highest MCC_Γ_, whereas the Dense structure yields the highest MCC_Γ_ for the binary and multinomial outcomes. Because the true support of Γ remains sparse in the Dense setting, these Γ metrics are well-defined. This contrast shows that a contemporaneous structure that is challenging for recovery Θ need not be challenging for recovering Γ, further illustrating the distinct behavior of the two model components.

**Table 2:**
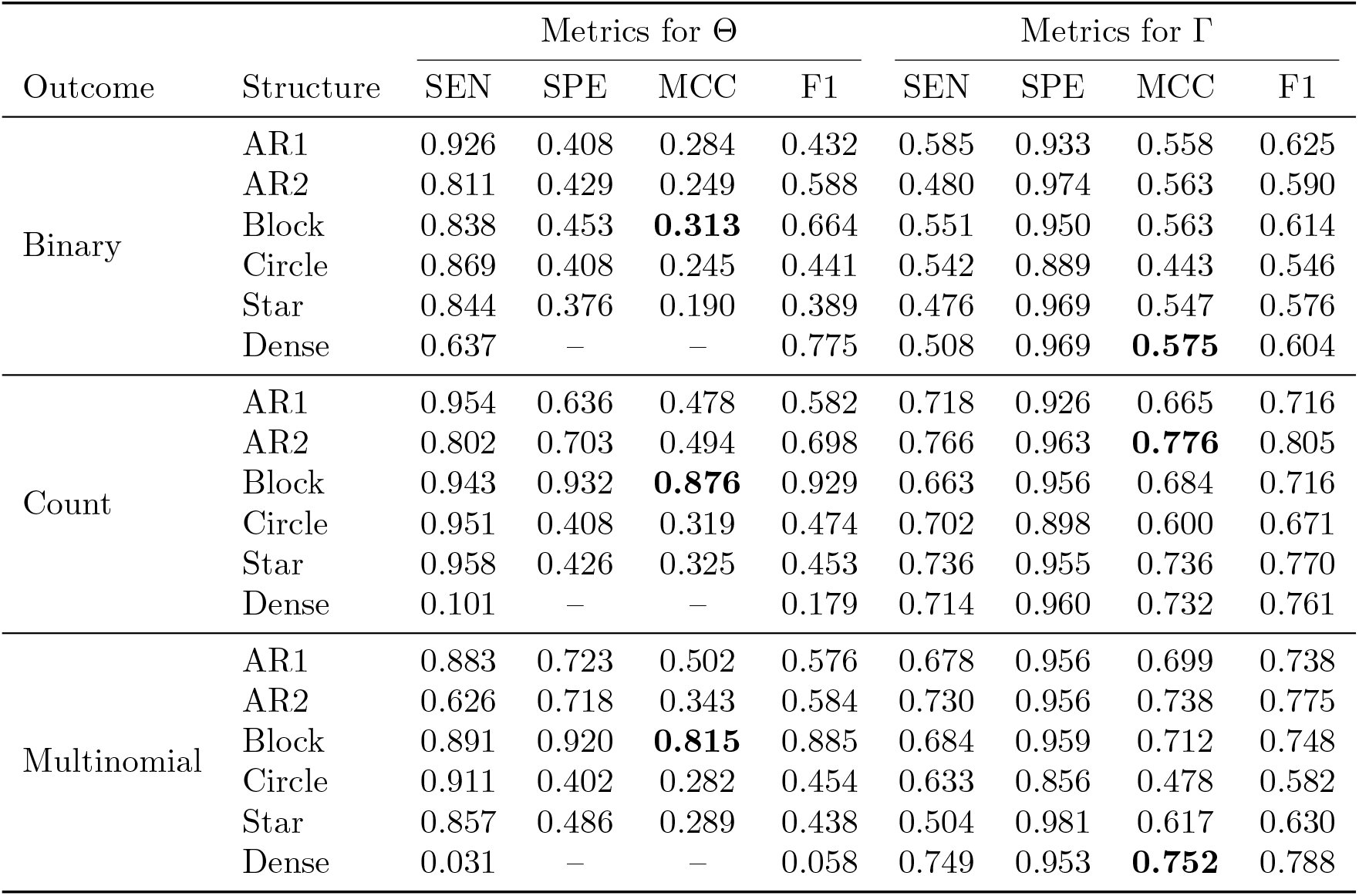
Support recovery under the baseline setting (*N, T, p, B*) = (100, 10, 10, 100). For the Dense structure, all off-diagonal elements of the true precision matrix are nonzero. Therefore, SPE and MCC are undefined for Θ and are denoted by “–”. All support recovery metrics remain well-defined for Γ, whose true support is sparse. Boldface indicates the numerically largest average MCC across the structures for which MCC is defined.

### 4.2 Effects of different factors

We next examine how support recovery changes as *N, T*, and *p* vary, while keeping the remaining parameters fixed at their baseline values. In the main text, we use MCC as the primary summary measure and focus on AR2, Block, and Star as representative contemporaneous dependence structures. These structures were chosen to represent distinct topological patterns for which MCC is well-defined. Full numerical results for SEN, SPE, MCC, and *F*_1_ score across all simulation settings are reported in the Supplementary Material.

Table 3 shows results beyond the baseline setting for three representative graph structures (AR2, Block and Star). For example, we see that increasing the sample size *N* improves recovery of Γ. Among the three representative structures, Block yields the numerically highest MCC_Θ_ for every sample size and outcome type. The improvement with *N* is clear for the count and multinomial outcomes, although the binary results are not strictly monotone. Thus, the favorable performance of Block for recovering Θ is maintained across the range of sample sizes considered. As for Γ, AR2 yields the highest MCC in most sample size settings. Overall, increasing *N* benefits both model components, while the structure associated with the best recovery differs between Θ and Γ.

**Table 3:**
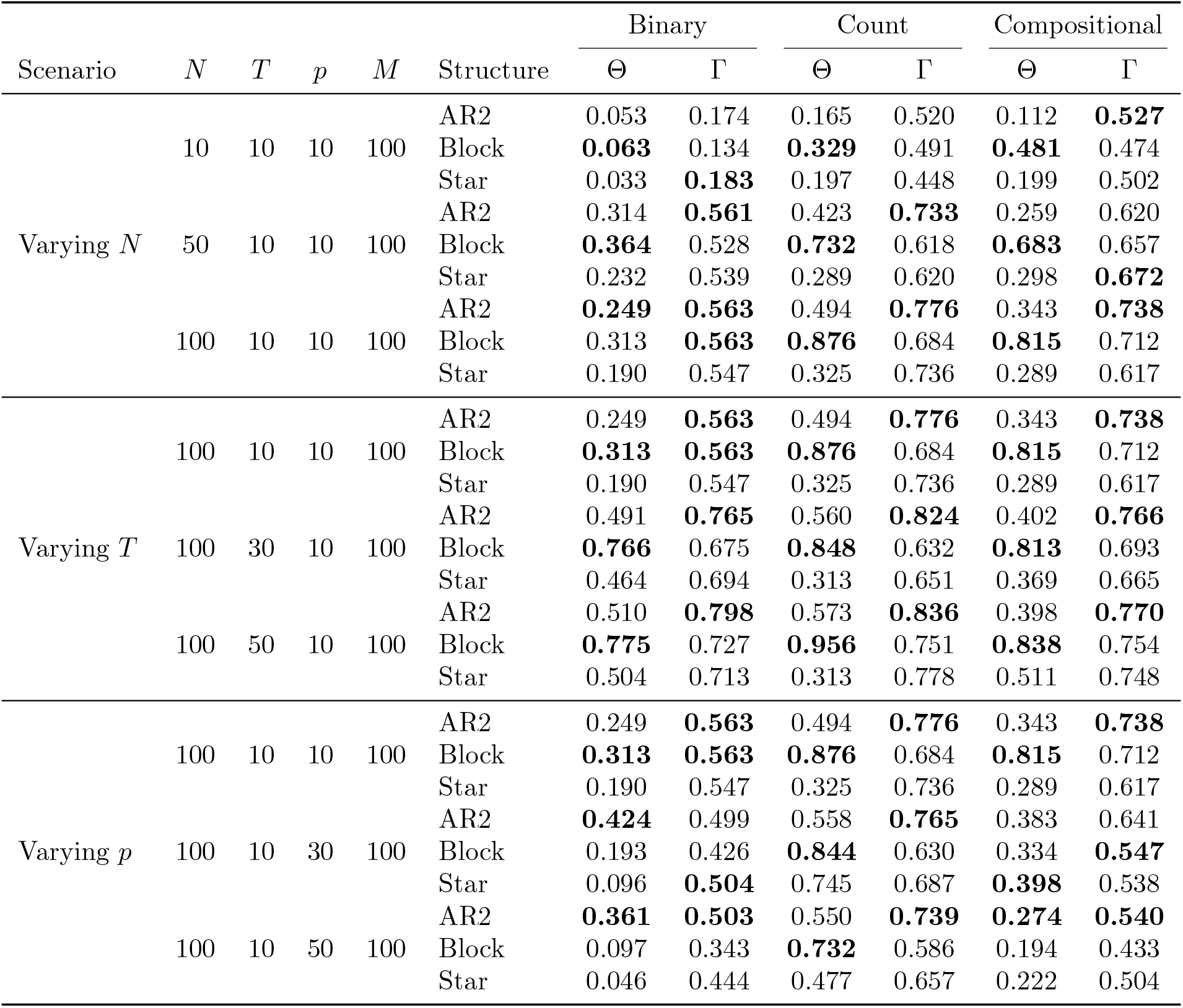
AR2, Block, and Star contemporaneous dependence structure results are shown below. Effect of sample size *N*, time length *T*, and latent dimension *p* on support recovery. Values are the average MCC over 100 simulation replicates. Boldface indicates the numerically largest average MCC across the structures for which MCC is defined. Complete results for all six structures are reported in the Supplementary Material.

| Scenario | $N$ | $T$ | $p$ | $M$ | Structure | Binary | | Count | | Compositional | |
| --- | --- | --- | --- | --- | --- | --- | --- | --- | --- | --- | --- |
| | | | | | | $\Theta$ | $\Gamma$ | $\Theta$ | $\Gamma$ | $\Theta$ | $\Gamma$ |
| Varying $N$ | 10 | 10 | 10 | 100 | AR2 | 0.053 | 0.174 | 0.165 | 0.520 | 0.112 | **0.527** |
|  |  |  |  |  | Block | **0.063** | 0.134 | **0.329** | 0.491 | **0.481** | 0.474 |
|  |  |  |  |  | Star | 0.033 | **0.183** | 0.197 | 0.448 | 0.199 | 0.502 |
|  | 50 | 10 | 10 | 100 | AR2 | 0.314 | **0.561** | 0.423 | **0.733** | 0.259 | 0.620 |
|  |  |  |  |  | Block | **0.364** | 0.528 | **0.732** | 0.618 | **0.683** | 0.657 |
|  |  |  |  |  | Star | 0.232 | 0.539 | 0.289 | 0.620 | 0.298 | **0.672** |
|  | 100 | 10 | 10 | 100 | AR2 | **0.249** | **0.563** | 0.494 | **0.776** | 0.343 | **0.738** |
|  |  |  |  |  | Block | 0.313 | **0.563** | **0.876** | 0.684 | **0.815** | 0.712 |
|  |  |  |  |  | Star | 0.190 | 0.547 | 0.325 | 0.736 | 0.289 | 0.617 |
| Varying $T$ | 100 | 10 | 10 | 100 | AR2 | 0.249 | **0.563** | 0.494 | **0.776** | 0.343 | **0.738** |
|  |  |  |  |  | Block | **0.313** | **0.563** | **0.876** | 0.684 | **0.815** | 0.712 |
|  |  |  |  |  | Star | 0.190 | 0.547 | 0.325 | 0.736 | 0.289 | 0.617 |
|  | 100 | 30 | 10 | 100 | AR2 | 0.491 | **0.765** | 0.560 | **0.824** | 0.402 | **0.766** |
|  |  |  |  |  | Block | **0.766** | 0.675 | **0.848** | 0.632 | **0.813** | 0.693 |
|  |  |  |  |  | Star | 0.464 | 0.694 | 0.313 | 0.651 | 0.369 | 0.665 |
|  | 100 | 50 | 10 | 100 | AR2 | 0.510 | **0.798** | 0.573 | **0.836** | 0.398 | **0.770** |
|  |  |  |  |  | Block | **0.775** | 0.727 | **0.956** | 0.751 | **0.838** | 0.754 |
|  |  |  |  |  | Star | 0.504 | 0.713 | 0.313 | 0.778 | 0.511 | 0.748 |
| Varying $p$ | 100 | 10 | 10 | 100 | AR2 | 0.249 | **0.563** | 0.494 | **0.776** | 0.343 | **0.738** |
|  |  |  |  |  | Block | **0.313** | **0.563** | **0.876** | 0.684 | **0.815** | 0.712 |
|  |  |  |  |  | Star | 0.190 | 0.547 | 0.325 | 0.736 | 0.289 | 0.617 |
|  | 100 | 10 | 30 | 100 | AR2 | **0.424** | 0.499 | 0.558 | **0.765** | 0.383 | 0.641 |
|  |  |  |  |  | Block | 0.193 | 0.426 | **0.844** | 0.630 | 0.334 | **0.547** |
|  |  |  |  |  | Star | 0.096 | **0.504** | 0.745 | 0.687 | **0.398** | 0.538 |
|  | 100 | 10 | 50 | 100 | AR2 | **0.361** | **0.503** | 0.550 | **0.739** | **0.274** | **0.540** |
|  |  |  |  |  | Block | 0.097 | 0.343 | **0.732** | 0.586 | 0.194 | 0.433 |
|  |  |  |  |  | Star | 0.046 | 0.444 | 0.477 | 0.657 | 0.222 | 0.504 |

Increasing the number of time points *T* generally improves recovery of Γ, because longer trajectories provide more observed transitions for estimating temporal dependence. Among the three displayed structures, AR2 yields the numerically highest MCC_Γ_ for all three outcome types at *T* = 30 and *T* = 50. A different pattern is observed for Θ. Block yields the highest MCC_Θ_ for every value of *T* and all three outcome types. The improvement with increasing *T* is pronounced for the binary outcome. For the count and multinomial outcomes, Block remains the strongest structure, although the increase is not strictly monotone at every intermediate value of *T* . These results reinforce the distinction between the two models’ components: the Block setting is favorable for recovering contemporaneous dependence, whereas the AR2 contemporaneous setting is generally favorable for recovering temporal dependence.

Increasing the latent dimension *p* makes support recovery more challenging, particularly for Γ, whose MCC tends to decrease as the number of parameters increases. AR2 yields the numerically highest MCC_Γ_ in most dimensional settings. For Θ, the structure with the highest MCC becomes less uniform as *p* increases. At the baseline dimension *p* = 10, Block yields the highest MCC_Θ_ for all three outcome types. For the count outcome, Block remains the strongest structure at *p* = 30 and *p* = 50. For the binary outcome, however, AR2 becomes numerically highest at both larger dimensions. For the multinomial outcome, Star is highest at *p* = 30, while AR2 is highest at *p* = 50. This shift indicates that the advantage of the Block structure for recovering Θ is strongest at low dimensions and is not uniform as *p* increases. In high-dimensional settings, recovery depends more on the graph topology and outcome type.

### 4.3 Outcome-specific trend

We also observe systematic differences across outcome types. Recovery under the binary observation model is more challenging, especially for the contemporaneous precision matrix Θ due to the fact that binary observations retain only binary information about the underlying latent state, whereas count and compositional observations provide additional count or categorical information. In these simulations, this difference is associated with stronger recovery under the count and multinomial models in many of the graphical settings.

Under the simulation settings considered, count outcomes generally yielded higher mean MCCs, particularly at larger dimensions, but this advantage was not uniform. For Θ under the Star structure at (*N, T, p*) = (100, 50, 10), compositional outcomes yielded a higher mean MCC than count outcomes (0.511 versus 0.313). Conversely, under the Block structure at (*N, T, p*) = (100, 10, 50), compositional outcomes yielded a lower mean MCC (0.194 versus 0.732), with low sensitivity and high specificity indicating missed true edges. Thus, neither observation model uniformly dominated, and weaker recovery involved different error patterns across settings. We therefore do not interpret the results as showing that one observation model uniformly dominates the other. Instead, the simulations indicate that recovery depends jointly on the observation model, graph structure, and dimension. In practice, the observation model should be selected according to the form of the observed data.

## 5. Integrative Human Microbiome Project (HMP2) Analysis

We applied the proposed method to longitudinal gut microbiome data from the HMP2 study, part of the Integrative Human Microbiome Project, which provides multi-omic profiles of host and microbial activity in inflammatory bowel disease (Lloyd-Price *and others*, 2019). The observations are compositional counts across microbial taxa, so we used the compositional observation model. The goal of this analysis was to estimate the lag-one temporal dependence structure, captured by Γ, and the contemporaneous conditional dependence network, captured by Θ, among dominant taxa.

We first restricted the HMP2 dataset to metagenomic taxonomic profiles and matched the taxonomic abundance table with the corresponding metadata using harmonized sample identifiers. To construct a balanced longitudinal dataset, we retained participants with samples from at least 10 distinct sampling weeks. Within each participant, we ordered samples by week and selected the first 10 distinct weeks. This procedure resulted in a balanced dataset containing *N* = 97 participants and *T* = 10 observations per participant, for a total of *NT* = 970 samples.

We selected taxa using these 970 samples. For each taxon, prevalence was defined as the proportion of samples in which the taxon had positive relative abundance, and mean relative abundance was calculated across all retained samples. We restricted the candidate features to species-level taxa with prevalence of at least 0.30 and positive mean relative abundance. Eligible taxa were ranked first by decreasing prevalence and then by decreasing mean relative abundance, and the top 11 taxa were selected for the multinomial analysis.

For each sample, the abundances of the selected 11 taxa were renormalized to sum to one. The resulting compositions were then converted to integer pseudo-counts with a fixed total of 100 per sample for use in the multinomial observation model. *Dorea longicatena* was used as the reference category, yielding a *p* = 10-dimensional latent representation. To assess the sensitivity of the feature selection procedure to the prevalence cutoff, we repeated the same ranking and selection procedure using thresholds of 0.10, 0.20, 0.30, and 0.50. The same 11 taxa were selected under all four thresholds, indicating that the retained features formed a stable set of dominant taxa in the balanced HMP2 cohort. We used the same compositional softmax observation model as described in Section 2.2.3. Briefly, for each subject *i* and time point *t*, the observed microbial composition was encoded as a compositional count vector

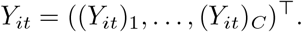

The latent state *Z*_*it*_ ∈ ℝ^*C*−1^ determined the category probabilities through the baseline category softmax link, with the *C*-th category used as the baseline. Conditional on *Z*_*it*_, *Y*_*it*_ followed a multinomial distribution with total count *M*_*it*_ and probability vector *π*_*it*_(*Z*_*it*_). For the HMP2 analysis, we selected *C* = 11 taxa, so the latent state dimension was *p* = *C* − 1 = 10. The selected relative abundance profiles were first normalized within the 11 selected taxa and then converted to fixed pseudo counts with *M*_*it*_ = 100, making the real data observations compatible with the compositional observation model.

We fitted the compositional dynamic graphical model over a two-dimensional grid of tuning parameters. The parameter *λ*_Γ_ controls the sparsity of the temporal transition matrix Γ, while *λ*_Θ_ controls the sparsity of the contemporaneous precision matrix Θ. For each candidate pair (*λ*_Γ_, *λ*_Θ_), the model was fitted using the proposed algorithm, producing estimates 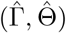. We then selected the final model as the one with the smallest EBIC value across the tuning grid. The selected tuning parameters were (*λ*_Γ_, *λ*_Θ_) = (0.30, 0.01), and the corresponding estimates 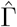 and 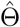 were used to construct the temporal and contemporaneous networks.

Figure 2 shows the estimated temporal and contemporaneous microbial association networks from the selected HMP2 data analysis. The temporal network estimated (Figure 2 (a)) from 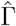 is sparse, indicating that only a small number of temporal cross-taxon associations were retained. In addition to these autoregressive effects, the temporal network identifies several directed cross-taxon associations. Under our plotting convention, a nonzero entry 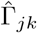 represents a directed edge from taxon *k* at time *t* − 1 to taxon *j* at time *t*. In particular, *F. prausnitzii* (F.prau in the figure) appears as a prominent node in the temporal network, with outgoing temporal edges to multiple Bacteroidetes taxa. These edges represent temporal conditional associations between the latent feature corresponding to *F. prausnitzii* at time *t* − 1 and the corresponding latent features of these taxa at time *t*, after accounting for the remaining latent features at time *t* − 1.

**Fig. 2:**
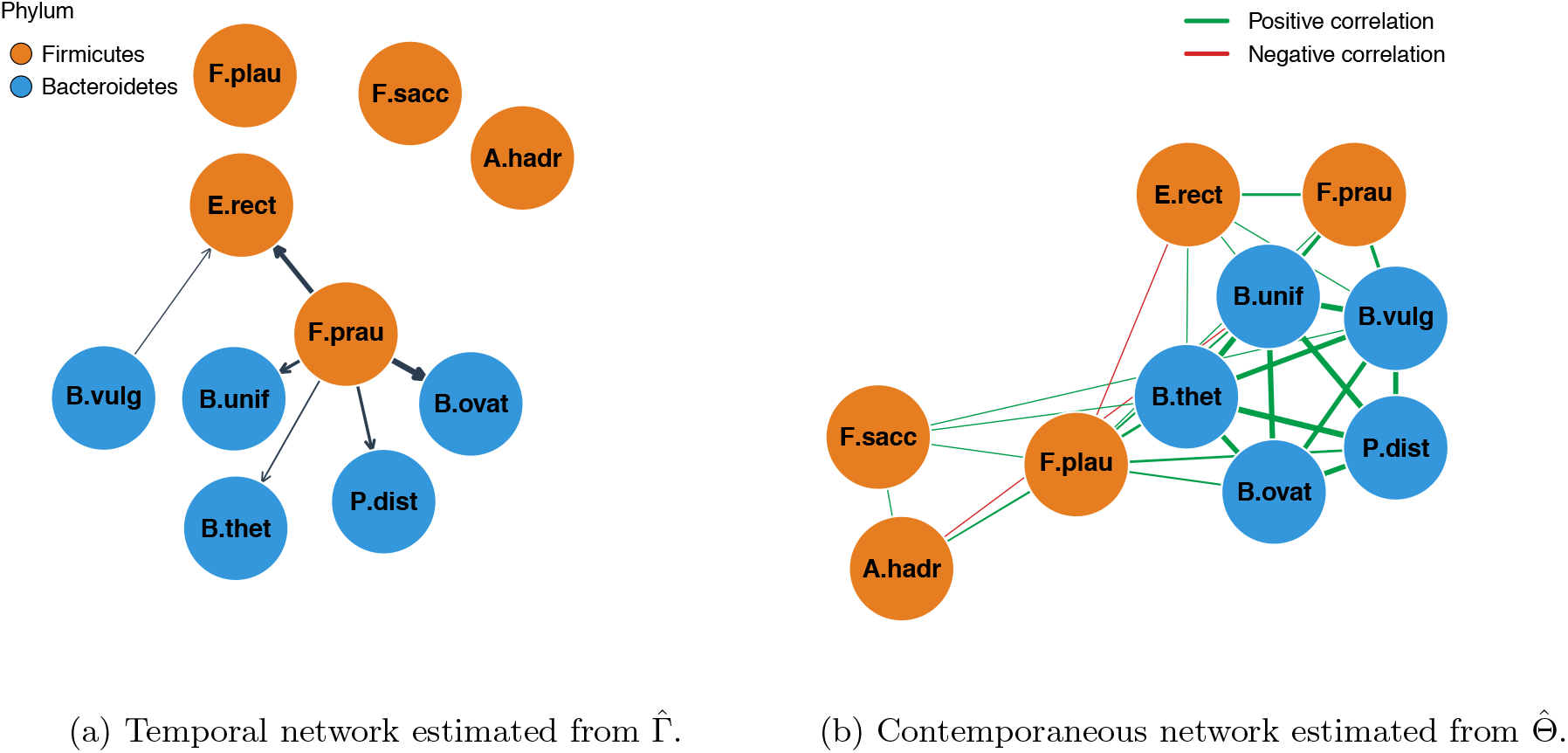
Estimated latent dynamic networks from the HMP2 longitudinal microbiome data (Lloyd-Price *and others*, 2019). Panel (a) displays the temporal network estimated from 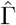. Nodes represent the *p* = *C ™* 1 = 10 non-baseline latent microbial components and are colored by phylum. Directed edges indicate nonzero lag-one effects, with edge direction determined by 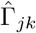 . Blue edges denote positive effects, and edge width is proportional to effect magnitude. Panel (b) displays the contemporaneous network estimated from 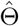. Nodes are colored by phylum. Undirected edges indicate nonzero off-diagonal entries of 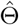, corresponding to conditional dependence relationships. For interpretability, edge colors and widths are determined by the induced contemporaneous correlation matrix from 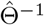: green indicates positive association, red indicates negative association, and edge width is proportional to effect magnitude.

The contemporaneous network (Figure 2 (b)) estimated from 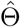 captures within-time conditional dependence among taxa. In addition, *F. prausnitzii* connects with multiple taxa in this cluster and with other Firmicutes taxa, suggesting that it may link taxa from different phylum-level groups in the estimated within-time association structure.

Overall, the HMP2 analysis illustrates the application of the proposed dynamic graphical model to longitudinal compositional microbiome data, yielding estimates of two complementary association networks. The estimated transition matrix 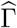 represents lag-one conditional associations among latent states, including taxon-specific persistence, while the estimated precision matrix 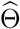 represents contemporaneous conditional dependence among latent innovations after accounting for lagged effects. Several taxa involved in the fitted networks have been highlighted in the HMP2 study and related microbiome research, providing biological context for the estimated associations. The HMP2 multi-omic association analysis identified *F. prausnitzii* as a highly connected taxonomic feature, with strong associations that were downregulated during dysbiosis (Lloyd-Price *and others*, 2019). More broadly, *F. prausnitzii* has been reported to have anti-inflammatory properties, and reduced abundance of this bacterium has been associated with inflammatory bowel disease (Sokol *and others*, 2008; Cao *and others*, 2014). In our fitted temporal network, outgoing edges from *F. prausnitzii* represent lag-one conditional associations between its corresponding latent state and those of other selected taxa, after accounting for the remaining latent states at the preceding time point. Its connections in the contemporaneous network represent conditional associations among the corresponding latent innovations.

The fitted temporal network also includes an association from *F. prausnitzii* to *P. distasonis*. The latter taxon was highlighted in the original HMP2 study through a nominal association with genetic variation at the *NKX2-3* locus, although this association did not remain significant after false discovery rate correction (Lloyd-Price *and others*, 2019). This previous host microbiome finding provides complementary biological context for *P. distasonis*, while addressing a different relationship from the temporal association estimated here. The fitted networks provide a description of temporal and contemporaneous dependence in the latent representation of microbiome compositions and generate hypotheses concerning associations among selected taxa for further biological investigation.

## 6. Discussion

### Findings

This paper proposes a latent Gaussian dynamic graphical model for longitudinal non-Gaussian responses, including binary, count, and compositional outcomes. The proposed framework jointly estimates two complementary dependence structures: the contemporaneous conditional dependence network encoded by Θ and the lag-one temporal dependence network encoded by Γ. While many existing dynamic graphical models are developed for Gaussian responses (Abegaz and Wit, 2013; Epskamp *and others*, 2018), and many non-Gaussian graphical models focus on cross-sectional dependence without explicitly estimating temporal transition effects (Yang *and others*, 2015; Lee and Hastie, 2015), our framework accommodates non-Gaussian observation models while jointly estimating both contemporaneous and temporal dependence.

Under the simulation settings considered here, the proposed method achieved varying degrees of support recovery for both Γ and Θ across the graph structures and outcome types examined. Recovery of the temporal dependence matrix Γ was generally more stable than recovery of the contemporaneous precision matrix Θ. This pattern may reflect differences in the two estimation tasks: estimation of Γ relies on temporal relationships between latent states at adjacent time points, whereas estimation of Θ requires recovering a sparse conditional dependence structure among the innovations and was more sensitive to the sparsity and topology of the underlying graph. In particular, recovery, especially for Θ, was more accurate for structured sparse graphs such as AR1 and Block than for the more challenging graph configurations considered.

Within these simulation settings, the binary observation model tended to yield lower support recovery performance. This may partly reflect the information about latent state variation conveyed by the simulated binary observations under the parameter configurations considered. The count and compositional observation models often yielded higher recovery, particularly for Θ, but neither observation model consistently outperformed the other across all settings. Their relative performance varied across graph structures, dimensions, and model components and, therefore, does not support a uniform ranking of the two observation models.

Over the ranges examined in the simulation study, increasing *N* or *T* and decreasing the latent dimension *p* generally improved recovery by increasing the available information or reducing the estimation burden. These improvements were more consistent for Γ, whereas recovery of Θ remained more sensitive to the sparsity and topology of the contemporaneous network.

In the HMP2 microbiome analysis, the selected taxa form a composition within each sample. After renormalizing their relative abundances and converting them to fixed total pseudo counts, we used a compositional observation model to represent the allocation of the total pseudo count across microbial categories, consistent with logistic normal multinomial approaches for microbiome compositional data (Xia *and others*, 2013). The fitted model separates two types of dependence. The transition matrix Γ captures lag-one temporal associations between taxa, while the precision matrix Θ captures contemporaneous conditional dependence after accounting for temporal dynamics. This distinction is useful because taxa may be associated at the same time point without having strong temporal predictive effects, and vice versa. Several directed cross-taxon associations are also identified, indicating potential temporal relationships among dominant taxa. The contemporaneous network is denser and reveals conditional associations among taxa within the same sampling time. In particular, several Bacteroidetes taxa form a connected cluster, and *F. prausnitzii* shows notable connections with multiple taxa. These findings are consistent with the known importance of *F. prausnitzii* in gut microbial ecology and inflammatory bowel disease (Lloyd-Price *and others*, 2019; Martín *and others*, 2023).

### Limitations and Future Work

There are several statistical limitations and directions for future work. The current implementation can be computationally demanding because each Monte Carlo iteration requires outcome-specific smoothing and Monte Carlo sampling of latent state trajectories for all subjects, and model selection requires repeated fitting over candidate penalty parameters. The computational burden increases with *N, T, p*, and the Monte Carlo sample size *M* . Independent simulation replicates were therefore distributed across the UW–Madison Center for High Throughput Computing. Improving the computational scalability of the estimation procedure is an important direction for future work. The proposed method relies on a Monte Carlo EM algorithm (Dempster *and others*, 1977; Wei and Tanner, 1990), and the Monte Carlo E-step can be computationally expensive as the dimensionality *p*, the time length *T*, or the number of Monte Carlo samples *B* increases. In addition, Monte Carlo variability may lead to small fluctuations in the convergence diagnostics after the initial decrease. Future work could improve computational efficiency through more scalable approximations, such as variational inference, stochastic approximation EM, adaptive Monte Carlo sampling, or more efficient Laplace-based approximations for the compositional model.

Next, tuning parameter selection is an important component of the proposed framework. In the simulation study, tuning parameters were selected using pilot runs and then fixed across 100 independent simulation replicates to improve computational feasibility and comparability across settings. While this design provides a practical way to summarize support recovery, it does not fully account for uncertainty in tuning parameter selection. Future work could consider more principled and data-adaptive tuning strategies, such as replicated EBIC selection (Chen and Chen, 2008; Foygel and Drton, 2010), predictive cross-validation, stability selection (Meinshausen and Bühlmann, 2010), or StARS-type procedures (Liu *and others*, 2010). It may also be useful to study the sensitivity of support recovery to the tuning grid to distinguish better procedures that choose different levels of sparsity for the contemporaneous and temporal components.

Finally, although the multinomial observation model is natural for the microbiome application considered here, further extensions would improve its flexibility for microbiome data. Microbiome sequencing data often exhibit zero inflation, overdispersion, compositional constraints, and potential covariate effects (Tang and Chen, 2019; Zeng *and others*, 2023). Extending the model to include subject-level and time-varying covariates, as well as zero-inflated or overdispersed observation models, would make the framework more broadly applicable. These extensions would also allow the estimated networks to better distinguish biological variation from technical and sampling variation in longitudinal microbiome studies.

## 7. ACKNOWLEDGMENTS

We thank Dr. Rosa Aghdam and Mason Garza for helpful discussions and feedback on earlier drafts of the manuscript.

## 10. Software availability

Code for the simulation experiments as well as R code that implement the approach are available on GitHub https://github.com/yyyyyyyyuhan/SDG-nongaussian.

