## Supplementary Material for "Sparse dynamic graphical models for longitudinal non-Gaussian data"

### List of Supplementary Algorithms

S1 Monte Carlo E-step for binary outcomes . . . . . 4
S2 Monte Carlo E-step for count outcomes . . . . . 4

### List of Supplementary Tables

S1 Full recovery results for the binary outcome when varying $N$. . . . . 10
S2 Full recovery results for the count outcome when varying $N$. . . . . 10
S3 Full recovery results for the compositional outcome when varying $N$. . . . . 11
S4 Full recovery results for the binary outcome when varying $T$. . . . . 11
S5 Full recovery results for the count outcome when varying $T$. . . . . 12
S6 Full recovery results for the compositional outcome when varying $T$. . . . . 12
S7 Full recovery results for the binary outcome when varying $p$. . . . . 13
S8 Full recovery results for the count outcome when varying $p$. . . . . 13
S9 Full recovery results for the compositional outcome when varying $p$. . . . . 14

This supplement provides additional methodological and numerical simulation details for the proposed estimation framework, which estimates the contemporaneous conditional dependence network and the lag-one temporal dependence structure. Section S1 presents the Monte Carlo E-step algorithms for binary and count outcomes. Section S2 shows the computational implementation details in the algorithm. Section S3 provides an additional matrix-level visualization of the contemporaneous dependence structures used in the simulation study. Section S4 reports the complete recovery results among all dependent structures, and Section S5 presents convergence diagnostics for selected pilot runs.

### S1. Additional Monte Carlo E-step algorithms

*Binary outcome.* For the binary probit model, the conditional expectations of the latent-state sufficient statistics in (2.22) are not available in closed form. We approximate these expectations using a Monte Carlo E-step based on latent Gaussian data augmentation (Albert and Chib, 1993; Holmes and Held, 2006). At iteration $k$, the Markov chain for subject $i$ is initialized at the final latent trajectory retained from the preceding iteration $Z_i^{(k-1)}$. Each Gibbs iteration first samples the components of the augmented variables $Z_i^*$, conditional on the current latent trajectory and the observed binary responses, from the truncated normal distributions in (2.4). Conditional on $Z_i^*$, the model is a linear Gaussian state-space model, and the complete latent trajectory is drawn using Gaussian simulation smoothing (Carter and Kohn, 1994; Durbin and Koopman, 2002). Conditioning on the current parameter estimates is suppressed here for notational simplicity.

For each subject, the sampler is run for $B + K$ iterations. The first $K$ draws are discarded as burn-in, and the cross-products of the temporal and current latent states are averaged over the remaining $B$ draws. The resulting subject-specific averages are then aggregated across subjects to form $S_{11}$, $S_{21}$, and $S_{22}$. The final latent trajectory is retained to initialize the sampler at the next iteration. Algorithm S1 gives the complete procedure.

*Count outcome.* For the count model, combining the Gaussian VAR(1) state equation in (2.1) with the lognormal-Poisson observation model in (2.5) yields a non-Gaussian smoothing distribution for the latent trajectory. Consequently, the conditional expectations of the latent-state sufficient statistics in (2.22) are not available in closed form. We approximate these expectations using an importance sampling smoother within the Monte Carlo E-step (Liu, 2001).

At each iteration and for each subject $i$, the smoother generates $B$ weighted latent trajectory draws,

$$\left\{ \left( Z_{i,1:T}^{(m)}, \widetilde{w}_{im} \right) \right\}_{m=1}^{B},$$

from a proposal distribution constructed for the count state-space model under the current parameter estimates. The unnormalized weights are converted to normalized weights,

$$w_{im} = \frac{\widetilde{w}_{im}}{\sum_{\ell=1}^{M} \widetilde{w}_{i\ell}}, \qquad \sum_{m=1}^{B} w_{im} = 1.$$

The subject contributions to $S_{11}$, $S_{21}$, and $S_{22}$ are then obtained from weighted averages of the corresponding latent-state quantities. These contributions are aggregated across subjects and used in the subsequent Monte Carlo M-step. Algorithm S2 summarizes the complete procedure.

**Algorithm S1:** Monte Carlo E-step for binary outcomes

**Input:** Observed binary trajectories $\boldsymbol{Y} = \{Y_{it}\}_{i=1,t=1}^{N,T}$; current parameters $(\Gamma^{(k-1)}, \Theta^{(k-1)})$; initial latent trajectories $\{Z_i^{(k-1)}\}_{i=1}^{N}$; retained Monte Carlo sample size $B$; burn-in $K$.

**Output:** Aggregated sufficient statistics $S_{11}$, $S_{21}$, $S_{22}$.

1 Initialize $S_{11} \leftarrow 0_{p\times p}$, $S_{21} \leftarrow 0_{p\times p}$, and $S_{22} \leftarrow 0_{p\times p}$; Set total number of iterations $L \leftarrow B + K$

2 **for** $i = 1, \ldots, N$ **do**

3     Initialize $Z_i^{\text{curr}} \leftarrow Z_i^{(k-1)}$; $E_{11} \leftarrow 0_{p\times p}$, $E_{21} \leftarrow 0_{p\times p}$, and $E_{22} \leftarrow 0_{p\times p}$

4     **for** $m = 1, \ldots, L$ **do**

5         For each $t = 1, \ldots, T$ and $j = 1, \ldots, p$, sample $(Z_{it}^*)_j$ from (2.4)

6         Draw

$$Z_i = (Z_{i1}, \ldots, Z_{iT}) \sim p\Big(Z_i \mid Z_i^*; \Gamma^{(k-1)}, \Theta^{(k-1)}\Big)$$

        using a Gaussian simulation smoother

7         Set $Z_i^{\text{curr}} \leftarrow Z_i$

8         **if** $m > K$ **then**

9             Let $Z_{i,1:(T-1)}$ and $Z_{i,2:T}$ be $(T-1)\times p$ matrices whose rows are $\{Z_{it}^\top\}_{t=1}^{T-1}$ and $\{Z_{it}^\top\}_{t=2}^{T}$, respectively

10             $E_{11} \leftarrow E_{11} + Z_{i,1:(T-1)}^\top Z_{i,1:(T-1)}$; $E_{21} \leftarrow E_{21} + Z_{i,2:T}^\top Z_{i,1:(T-1)}$; $E_{22} \leftarrow E_{22} + Z_{i,2:T}^\top Z_{i,2:T}$.

11     Set $Z_i^{(k)} \leftarrow Z_i^{\text{curr}}$; $S_{11} \leftarrow S_{11} + E_{11}/B$; $S_{21} \leftarrow S_{21} + E_{21}/B$; $S_{22} \leftarrow S_{22} + E_{22}/B$.

12 Symmetrize $S_{11} \leftarrow (S_{11} + S_{11}^\top)/2$ and $S_{22} \leftarrow (S_{22} + S_{22}^\top)/2$.

13 **return** $S_{11}, S_{21}, S_{22}$; $\{Z_i^{(k)}\}_{i=1}^{N}$.

**Algorithm S2:** Monte Carlo E-step for count outcomes

**Input:** Observed count trajectories $\boldsymbol{Y} = \{Y_{it}\}_{i=1,t=1}^{N,T}$; current parameters $(\Gamma^{(k-1)}, \Theta^{(k-1)})$; Monte Carlo sample size $B$.

**Output:** Aggregated sufficient statistics $S_{11}$, $S_{21}$, and $S_{22}$.

1 Initialize $S_{11} \leftarrow 0_{p\times p}$, $S_{21} \leftarrow 0_{p\times p}$, and $S_{22} \leftarrow 0_{p\times p}$

2 **for** $i = 1, \ldots, N$ **do**

3     Apply an importance sampling smoother to the count state-space model for subject $i$ under $(\Gamma^{(k-1)}, \Theta^{(k-1)})$, obtaining weighted latent trajectory draws $\{(Z_{i,1:T}^{(m)}, \widetilde{w}_{im})\}_{m=1}^{B}$

4     Set $w_{im} \leftarrow \frac{\widetilde{w}_{im}}{\sum_{\ell=1}^{B} \widetilde{w}_{i\ell}}, \qquad m = 1, \ldots, B.$

5     Initialize $E_{11} \leftarrow 0_{p\times p}$, $E_{21} \leftarrow 0_{p\times p}$, and $E_{22} \leftarrow 0_{p\times p}$

6     **for** $m = 1, \ldots, B$ **do**

7         Let $Z_{i,1:(T-1)}^{(m)}$ and $Z_{i,2:T}^{(m)}$ be $(T-1)\times p$ matrices whose rows are $\{(Z_{it}^{(m)})^\top\}_{t=1}^{T-1}$ and $\{(Z_{it}^{(m)})^\top\}_{t=2}^{T}$, respectively

8         $E_{11} \leftarrow E_{11} + w_{im}(Z_{i,1:(T-1)}^{(m)})^\top Z_{i,1:(T-1)}^{(m)}$; $E_{21} \leftarrow E_{21} + w_{im}(Z_{i,2:T}^{(m)})^\top Z_{i,1:(T-1)}^{(m)}$; $E_{22} \leftarrow E_{22} + w_{im}(Z_{i,2:T}^{(m)})^\top Z_{i,2:T}^{(m)}$

9     $S_{11} \leftarrow S_{11} + E_{11}$; $S_{21} \leftarrow S_{21} + E_{21}$; $S_{22} \leftarrow S_{22} + E_{22}$;

10 Symmetrize $S_{11} \leftarrow (S_{11} + S_{11}^\top)/2$ and $S_{22} \leftarrow (S_{22} + S_{22}^\top)/2$

11 **return** $S_{11},\ S_{21},\ S_{22}$

### S2. Computational implementation

The three outcome-specific implementations share the same two-stage penalized update, but use different Monte Carlo E-steps. State-space filtering, smoothing, and simulation are implemented using the R package `KFAS` (Helske, 2017). At each iteration, the outcome-specific E-step produces approximations of the same complete data sufficient statistics, $S_{11}$, $S_{21}$, and $S_{22}$. These statistics are subsequently passed to a common update for the transition matrix $\Gamma$ and the innovation precision matrix $\Theta$.

*Binary E-step.* For binary outcomes, we combine probit data augmentation with Gaussian simulation smoothing. Conditional on the current latent trajectory, the augmented observations are generated componentwise from truncated normal distributions. Conditional on these augmented observations, a complete latent trajectory is drawn with `KFAS::simulateSSM`. For each subject, the sampler is run for $B+M$ iterations, the first $B$ draws are discarded, and the sufficient statistics are averaged over the remaining $M$ trajectories. The final latent trajectory is retained to initialize the Markov chain at the next iteration. Full details are given in Algorithm S1.

*Count E-step.* For count outcomes, the observation model is specified in `KFAS` using its Poisson state-space functionality. For each subject, weighted latent trajectories are generated using `KFAS::importanceSSM` with antithetic sampling. The returned importance weights are restricted to finite, nonnegative values and normalized to sum to one. If the returned weights cannot be normalized, uniform weights are used as a numerical fallback. The normalized weights are then used to construct weighted estimates of $S_{11}$, $S_{21}$, and $S_{22}$.

*Composition E-step.* For composition outcomes, `KFAS` does not directly provide a multinomial observation model, we construct a Laplace-based pseudo-Gaussian state-space approximation as in (2.15). The pseudo-observation and covariance matrix define a Gaussian state-space model that is filtered and smoothed using `KFAS::KFS`. After constructing the Gaussian approximation, $M$ latent trajectories are generated using `KFAS::simulateSSM`. The importance weights are computed on the log scale as the difference between the exact compositional log likelihood and the pseudo-Gaussian observation log likelihood. The resulting weighted trajectories are used to construct $S_{11}$, $S_{21}$, and $S_{22}$, and their weighted mean is used to initialize the Laplace approximation at the iteration.

*Common M-step.* Since the sufficient statistics are aggregated over all observed transitions, the working loss for $\Gamma$ is normalized by $N_{\text{eff}} = N(T-1)$. The transition matrix is updated using a coordinate-gradient descent solver for the SCAD penalized objective in Algorithm 2, with SCAD parameter $a = 3.7$ (Fan and Li, 2001). The estimates from the preceding iteration is used to initialize the coordinate solver. Given the updated $\Gamma$, the innovation precision matrix $\Theta$ is estimated from the residual covariance matrix using the `glasso` package using $\ell_1$ penalty is applied to all entries of $\Theta$ (Friedman *and others*, 2008, 2019). The estimated contemporaneous graph is nevertheless defined only through the off-diagonal support of $\Theta$.

*Numerical stabilization and iteration control.* Covariance matrices were symmetrized and augmented with a diagonal ridge term of $10^{-6}I$ as numerical safeguards against numerical asymmetry and near-singularity. This fixed numerical adjustment was distinct from the sparsity inducing penalties used to estimate $\Gamma$ and $\Theta$.

Unless user-specified initial values were supplied, the binary and count implementations initialized the transition matrix at $\Gamma^{(0)} = 0.5I$, whereas the compositional implementation used $\Gamma^{(0)} = 0.2I$. Both choices have spectral radii strictly smaller than one and therefore correspond to stable initial VAR(1) transition matrices. All three implementations initialized $\Theta^{(0)} = I$, providing a positive definite starting value with no off-diagonal conditional associations. These default matrices served as starting values rather than constraints on subsequent parameter estimates.

During the first five iterations, both penalty parameters were temporarily set to 0.05. The implementation then switched to the selected values $(\lambda_\Gamma, \lambda_\Theta)$. This preliminary stage provided starting estimates for subsequent iterations using the selected penalties. The initial penalty value and the duration of this stage were fixed implementation choices rather than additional data-dependent tuning parameters.

To monitor stabilization of the parameter estimates, we recorded the relative Frobenius changes at each iteration:

$$\Delta_\Gamma^{(k)} = \frac{\|\Gamma^{(k)} - \Gamma^{(k-1)}\|_F}{\|\Gamma^{(k-1)}\|_F + 10^{-10}}, \qquad \Delta_\Theta^{(k)} = \frac{\|\Theta^{(k)} - \Theta^{(k-1)}\|_F}{\|\Theta^{(k-1)}\|_F + 10^{-10}}.$$

The constant $10^{-10}$ prevents division by zero when the previous iterate has zero Frobenius norm. These quantities were recorded for all three outcome models.

### S3. Additional visualization of simulation structures

The simulation study considers six contemporaneous dependence structures: AR1, AR2, Block, Circle, Star, and Dense. Figure 1 in the main text provides graphical representations of these structures. Here, we provide an additional matrix-level visualization of the six contemporaneous dependence structures used in the simulation study. Figure S1 shows the numerical values of the matrices used to construct the corresponding settings when $p = 10$. The AR1 and Block settings were constructed by specifying a covariance matrix $\Sigma$; the corresponding panels therefore display $\Sigma$, with the true precision matrix defined as $\Theta = \Sigma^{-1}$. The AR2, Circle, Star, and Dense settings were constructed by specifying $\Theta$, and their panels consequently display the precision matrix. In all six settings, support recovery is evaluated with respect to the off-diagonal support of the resulting true precision matrix $\Theta$.

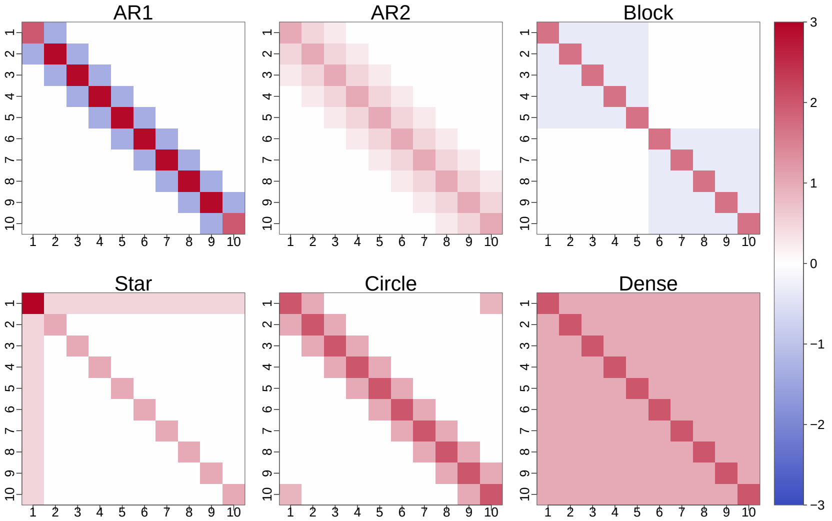

Fig. S1: Heatmaps of the true precision matrices $\Theta$ for the six contemporaneous dependence structures used in the simulation study at $p = 10$.

### S4. Complete simulation results

Tables S1–S9 provide the complete support recovery, summarizing the underlying six contemporaneous dependence structures in the main text. Tables S1–S3 examine the effect of sample size $N$, Tables S4–S6 examine the effect of time-series length $T$, and Tables S7–S9 examine the effect of latent dimension $p$. For each factor, separate tables are provided for the binary, count, and compositional outcomes. All six contemporaneous dependence structures are included, and SEN, SPE, MCC, and $F_1$ are reported for both $\Theta$ and $\Gamma$. All entries are averages over 100 simulation replicates. When one factor is varied, the remaining parameters are fixed at their baseline values. Specifically, $(T, p, B) = (10, 10, 100)$ when $N$ is varied, $(N, p, B) = (100, 10, 100)$ when $T$ is varied, and $(N, T, B) = (100, 10, 100)$ when $p$ is varied. Within each parameter setting and target matrix, boldface indicates the largest MCC across the graph structures for which MCC is defined. For the Dense structure, all off-diagonal entries of the true $\Theta$ are nonzero. Consequently, SPE and MCC are undefined for $\Theta$ and are denoted by "–". All four metrics remain well defined for $\Gamma$, whose true support is sparse.

#### *S4.1 Effect of sample size N*

Tables S1–S3 show that recovery of $\Gamma$ improves markedly from $N = 10$ to $N = 50$ across all three outcome types and all six contemporaneous dependence structures. Improvements from $N = 50$ to $N = 100$ are smaller and are not always monotone. Recovery of $\Theta$ is more dependent on the outcome type and graph structure. Under the count and compositional outcomes, the improvement is pronounced for the Block structure. Thus, increasing $N$ benefits recovery of $\Gamma$ more consistently than recovery of $\Theta$.

#### *S4.2 Effect of time-series length T*

Tables S4–S6 show that longer time series generally improve the recovery of $\Gamma$, with a particularly consistent improvement under AR2 across the three outcome types. Recovery of $\Theta$ also improves in several settings, most notably for the binary outcome under the Block structure. For the count and compositional outcomes, however, the changes in $\mathrm{MCC}_\Theta$ are smaller and are not uniformly monotone across structures. These results indicate that increasing $T$ primarily benefits the estimation of lagged dependence, while its effect on contemporaneous dependence remains structure-dependent.

#### *S4.3 Effect of latent dimension p*

Tables S7–S9 show that recovery of $\Gamma$ generally becomes more difficult as $p$ increases, with MCC at $p = 50$ typically lower than at $p = 10$. For $\Theta$, the effect is again structure-dependent. Deterioration is most apparent under the Block structure, whereas AR1 and Circle exhibits less uniform patterns. In several higher-dimensional settings, increased specificity is accompanied by reduced sensitivity, indicating more conservative edge selection rather than uniform improvement in support recovery. Overall, the effect of dimension on $\Theta$-recovery cannot be characterized independently of graph structure.

Table S1: Full recovery results for the binary outcome when varying $N$. Values are averaged over 100 simulation replicates. Boldface indicates the numerically largest average MCC across the structures for which MCC is defined.

| $N$ | $T$ | $p$ | $B$ | Rep. | Structure | Metrics for $\Theta$ | | | | Metrics for $\Gamma$ | | | |
|---|---|---|---|---|---|---|---|---|---|---|---|---|---|
| | | | | | | SEN | SPE | MCC | F1 | SEN | SPE | MCC | F1 |
| 100 | 10 | 10 | 100 | 100 | AR1 | 0.926 | 0.408 | 0.284 | 0.432 | 0.585 | 0.933 | 0.558 | 0.625 |
| | | | | | AR2 | 0.811 | 0.429 | 0.249 | 0.588 | 0.480 | 0.974 | 0.563 | 0.590 |
| | | | | | Block | 0.838 | 0.453 | **0.313** | 0.664 | 0.551 | 0.950 | 0.563 | 0.614 |
| | | | | | Circle | 0.869 | 0.408 | 0.245 | 0.441 | 0.542 | 0.889 | 0.443 | 0.546 |
| | | | | | Star | 0.844 | 0.376 | 0.190 | 0.389 | 0.476 | 0.969 | 0.547 | 0.576 |
| | | | | | Dense | 0.637 | – | – | 0.775 | 0.508 | 0.969 | **0.575** | 0.604 |
| 50 | 10 | 10 | 100 | 100 | AR1 | 0.800 | 0.673 | **0.387** | 0.518 | 0.679 | 0.811 | 0.437 | 0.557 |
| | | | | | AR2 | 0.596 | 0.720 | 0.314 | 0.573 | 0.694 | 0.889 | **0.561** | 0.648 |
| | | | | | Block | 0.626 | 0.733 | 0.364 | 0.636 | 0.625 | 0.900 | 0.528 | 0.615 |
| | | | | | Circle | 0.762 | 0.597 | 0.301 | 0.480 | 0.779 | 0.445 | 0.185 | 0.395 |
| | | | | | Star | 0.628 | 0.655 | 0.232 | 0.417 | 0.605 | 0.917 | 0.539 | 0.619 |
| | | | | | Dense | 0.338 | – | – | 0.501 | 0.553 | 0.946 | 0.559 | 0.615 |
| 10 | 10 | 10 | 100 | 100 | AR1 | 0.678 | 0.406 | 0.070 | 0.333 | 0.648 | 0.514 | 0.133 | 0.362 |
| | | | | | AR2 | 0.632 | 0.421 | 0.053 | 0.486 | 0.739 | 0.472 | 0.174 | 0.385 |
| | | | | | Block | 0.648 | 0.414 | 0.063 | 0.543 | 0.736 | 0.426 | 0.134 | 0.366 |
| | | | | | Circle | 0.635 | 0.472 | **0.091** | 0.363 | 0.886 | 0.152 | 0.036 | 0.337 |
| | | | | | Star | 0.630 | 0.411 | 0.033 | 0.316 | 0.678 | 0.545 | 0.183 | 0.389 |
| | | | | | Dense | 0.590 | – | – | 0.740 | 0.649 | 0.605 | **0.208** | 0.403 |

Table S2: Full recovery results for the count outcome when varying $N$. Values are averaged over 100 simulation replicates. Boldface indicates the numerically largest average MCC across the structures for which MCC is defined.

| $N$ | $T$ | $p$ | $B$ | Rep. | Structure | Metrics for $\Theta$ | | | | Metrics for $\Gamma$ | | | |
|---|---|---|---|---|---|---|---|---|---|---|---|---|---|
| | | | | | | SEN | SPE | MCC | F1 | SEN | SPE | MCC | F1 |
| 100 | 10 | 10 | 100 | 100 | AR1 | 0.954 | 0.636 | 0.478 | 0.562 | 0.718 | 0.926 | 0.665 | 0.716 |
| | | | | | AR2 | 0.802 | 0.703 | 0.494 | 0.698 | 0.766 | 0.963 | **0.776** | 0.805 |
| | | | | | Block | 0.943 | 0.932 | **0.876** | 0.929 | 0.663 | 0.956 | 0.684 | 0.716 |
| | | | | | Circle | 0.951 | 0.408 | 0.319 | 0.474 | 0.702 | 0.898 | 0.600 | 0.671 |
| | | | | | Star | 0.958 | 0.426 | 0.325 | 0.453 | 0.736 | 0.955 | 0.736 | 0.770 |
| | | | | | Dense | 0.101 | – | – | 0.179 | 0.714 | 0.960 | 0.732 | 0.761 |
| 50 | 10 | 10 | 100 | 100 | AR1 | 0.978 | 0.568 | 0.441 | 0.530 | 0.598 | 0.955 | 0.627 | 0.661 |
| | | | | | AR2 | 0.789 | 0.642 | 0.423 | 0.662 | 0.730 | 0.957 | **0.733** | 0.769 |
| | | | | | Block | 0.910 | 0.822 | **0.732** | 0.854 | 0.545 | 0.970 | 0.618 | 0.639 |
| | | | | | Circle | 0.937 | 0.410 | 0.308 | 0.469 | 0.675 | 0.888 | 0.561 | 0.639 |
| | | | | | Star | 0.908 | 0.438 | 0.289 | 0.439 | 0.533 | 0.975 | 0.620 | 0.634 |
| | | | | | Dense | 0.200 | – | – | 0.328 | 0.595 | 0.970 | 0.657 | 0.680 |
| 10 | 10 | 10 | 100 | 100 | AR1 | 0.848 | 0.576 | **0.346** | 0.482 | 0.495 | 0.918 | 0.458 | 0.529 |
| | | | | | AR2 | 0.751 | 0.409 | 0.165 | 0.550 | 0.619 | 0.893 | **0.520** | 0.604 |
| | | | | | Block | 0.773 | 0.549 | 0.329 | 0.660 | 0.481 | 0.938 | 0.491 | 0.541 |
| | | | | | Circle | 0.883 | 0.358 | 0.219 | 0.428 | 0.632 | 0.677 | 0.263 | 0.433 |
| | | | | | Star | 0.791 | 0.445 | 0.197 | 0.395 | 0.669 | 0.817 | 0.448 | 0.558 |
| | | | | | Dense | 0.469 | – | – | 0.632 | 0.492 | 0.945 | 0.514 | 0.560 |

Table S3: Full recovery results for the compositional outcome when varying $N$. Values are averaged over 100 simulation replicates. Boldface indicates the numerically largest average MCC across the structures for which MCC is defined.

| $N$ | $T$ | $p$ | $B$ | Rep. | Structure | Metrics for $\Theta$ | | | | Metrics for $\Gamma$ | | | |
|---|---|---|---|---|---|---|---|---|---|---|---|---|---|
| | | | | | | SEN | SPE | MCC | F1 | SEN | SPE | MCC | F1 |
| 100 | 10 | 10 | 100 | 100 | AR1 | 0.883 | 0.723 | 0.502 | 0.576 | 0.678 | 0.956 | 0.699 | 0.738 |
| | | | | | AR2 | 0.626 | 0.718 | 0.343 | 0.584 | 0.730 | 0.956 | 0.738 | 0.775 |
| | | | | | Block | 0.891 | 0.920 | **0.815** | 0.885 | 0.684 | 0.959 | 0.712 | 0.748 |
| | | | | | Circle | 0.911 | 0.402 | 0.282 | 0.454 | 0.633 | 0.856 | 0.478 | 0.582 |
| | | | | | Star | 0.857 | 0.486 | 0.289 | 0.438 | 0.504 | 0.981 | 0.617 | 0.630 |
| | | | | | Dense | 0.031 | – | – | 0.058 | 0.749 | 0.953 | **0.752** | 0.788 |
| 50 | 10 | 10 | 100 | 100 | AR1 | 0.842 | 0.759 | 0.512 | 0.588 | 0.578 | 0.965 | 0.637 | 0.669 |
| | | | | | AR2 | 0.662 | 0.599 | 0.259 | 0.559 | 0.509 | 0.982 | 0.620 | 0.636 |
| | | | | | Block | 0.867 | 0.816 | **0.683** | 0.819 | 0.590 | 0.969 | 0.657 | 0.686 |
| | | | | | Circle | 0.890 | 0.390 | 0.255 | 0.439 | 0.567 | 0.867 | 0.434 | 0.544 |
| | | | | | Star | 0.862 | 0.494 | 0.298 | 0.439 | 0.672 | 0.945 | 0.672 | 0.719 |
| | | | | | Dense | 0.126 | – | – | 0.219 | 0.638 | 0.968 | **0.695** | 0.724 |
| 10 | 10 | 10 | 100 | 100 | AR1 | 0.761 | 0.733 | 0.417 | 0.521 | 0.602 | 0.877 | 0.486 | 0.580 |
| | | | | | AR2 | 0.698 | 0.410 | 0.112 | 0.521 | 0.517 | 0.942 | 0.527 | 0.588 |
| | | | | | Block | 0.596 | 0.862 | **0.481** | 0.644 | 0.662 | 0.836 | 0.474 | 0.582 |
| | | | | | Circle | 0.774 | 0.365 | 0.124 | 0.385 | 0.468 | 0.805 | 0.258 | 0.416 |
| | | | | | Star | 0.347 | 0.846 | 0.199 | 0.336 | 0.451 | 0.956 | 0.502 | 0.545 |
| | | | | | Dense | 0.303 | – | – | 0.458 | 0.477 | 0.960 | **0.536** | 0.573 |

Table S4: Full recovery results for the binary outcome when varying $T$. Values are averaged over 100 simulation replicates. Boldface indicates the numerically largest average MCC across the structures for which MCC is defined.

| $N$ | $T$ | $p$ | $B$ | Rep. | Structure | Metrics for $\Theta$ | | | | Metrics for $\Gamma$ | | | |
|---|---|---|---|---|---|---|---|---|---|---|---|---|---|
| | | | | | | SEN | SPE | MCC | F1 | SEN | SPE | MCC | F1 |
| 100 | 10 | 10 | 100 | 100 | AR1 | 0.926 | 0.408 | 0.284 | 0.432 | 0.585 | 0.933 | 0.558 | 0.625 |
| | | | | | AR2 | 0.811 | 0.429 | 0.249 | 0.588 | 0.480 | 0.974 | 0.563 | 0.590 |
| | | | | | Block | 0.838 | 0.453 | **0.313** | 0.664 | 0.551 | 0.950 | 0.563 | 0.614 |
| | | | | | Circle | 0.869 | 0.408 | 0.245 | 0.441 | 0.542 | 0.889 | 0.443 | 0.546 |
| | | | | | Star | 0.844 | 0.376 | 0.190 | 0.389 | 0.476 | 0.969 | 0.547 | 0.576 |
| | | | | | Dense | 0.637 | – | – | 0.775 | 0.508 | 0.969 | **0.575** | 0.604 |
| 100 | 30 | 10 | 100 | 100 | AR1 | 0.917 | 0.779 | 0.585 | 0.658 | 0.736 | 0.933 | 0.675 | 0.735 |
| | | | | | AR2 | 0.608 | 0.864 | 0.491 | 0.652 | 0.733 | 0.976 | **0.765** | 0.799 |
| | | | | | Block | 0.803 | 0.946 | **0.766** | 0.852 | 0.773 | 0.919 | 0.675 | 0.738 |
| | | | | | Circle | 0.862 | 0.646 | 0.426 | 0.557 | 0.669 | 0.868 | 0.508 | 0.608 |
| | | | | | Star | 0.776 | 0.769 | 0.464 | 0.576 | 0.650 | 0.973 | 0.694 | 0.732 |
| | | | | | Dense | 0.138 | – | – | 0.238 | 0.726 | 0.965 | 0.737 | 0.776 |
| 100 | 50 | 10 | 100 | 100 | AR1 | 0.920 | 0.775 | 0.583 | 0.656 | 0.680 | 0.960 | 0.690 | 0.736 |
| | | | | | AR2 | 0.610 | 0.876 | 0.510 | 0.658 | 0.800 | 0.970 | **0.798** | 0.832 |
| | | | | | Block | 0.814 | 0.944 | **0.775** | 0.860 | 0.718 | 0.964 | 0.727 | 0.769 |
| | | | | | Circle | 0.867 | 0.650 | 0.435 | 0.564 | 0.646 | 0.912 | 0.562 | 0.645 |
| | | | | | Star | 0.802 | 0.785 | 0.504 | 0.605 | 0.667 | 0.975 | 0.713 | 0.749 |
| | | | | | Dense | 0.132 | – | – | 0.228 | 0.645 | 0.980 | 0.711 | 0.739 |

Table S5: Full recovery results for the count outcome when varying $T$. Values are averaged over 100 simulation replicates. Boldface indicates the numerically largest average MCC across the structures for which MCC is defined.

| $N$ | $T$ | $p$ | $B$ | Rep. | Structure | Metrics for $\Theta$ | | | | Metrics for $\Gamma$ | | | |
|---|---|---|---|---|---|---|---|---|---|---|---|---|---|
| | | | | | | SEN | SPE | MCC | F1 | SEN | SPE | MCC | F1 |
| 100 | 10 | 10 | 100 | 100 | AR1 | 0.954 | 0.636 | 0.478 | 0.562 | 0.718 | 0.926 | 0.665 | 0.716 |
| | | | | | AR2 | 0.802 | 0.703 | 0.494 | 0.698 | 0.766 | 0.963 | **0.776** | 0.805 |
| | | | | | Block | 0.943 | 0.932 | **0.876** | 0.929 | 0.663 | 0.956 | 0.684 | 0.716 |
| | | | | | Circle | 0.951 | 0.408 | 0.319 | 0.474 | 0.702 | 0.898 | 0.600 | 0.671 |
| | | | | | Star | 0.958 | 0.426 | 0.325 | 0.453 | 0.736 | 0.955 | 0.736 | 0.770 |
| | | | | | Dense | 0.101 | – | – | 0.179 | 0.714 | 0.960 | 0.732 | 0.761 |
| 100 | 30 | 10 | 100 | 100 | AR1 | 0.960 | 0.646 | 0.489 | 0.570 | 0.652 | 0.963 | 0.688 | 0.715 |
| | | | | | AR2 | 0.820 | 0.753 | 0.560 | 0.735 | 0.827 | 0.964 | 0.824 | 0.849 |
| | | | | | Block | 0.935 | 0.913 | **0.848** | 0.915 | 0.534 | 0.981 | 0.632 | 0.642 |
| | | | | | Circle | 0.964 | 0.409 | 0.333 | 0.479 | 0.805 | 0.854 | 0.627 | 0.698 |
| | | | | | Star | 0.954 | 0.414 | 0.313 | 0.447 | 0.551 | 0.983 | 0.651 | 0.661 |
| | | | | | Dense | 0.040 | – | – | 0.075 | 0.703 | 0.977 | **0.757** | 0.776 |
| 100 | 50 | 10 | 100 | 100 | AR1 | 0.959 | 0.643 | 0.485 | 0.566 | 0.644 | 0.963 | 0.682 | 0.709 |
| | | | | | AR2 | 0.827 | 0.759 | 0.573 | 0.741 | 0.826 | 0.971 | **0.836** | 0.858 |
| | | | | | Block | 0.966 | 0.987 | **0.956** | 0.973 | 0.710 | 0.972 | 0.751 | 0.773 |
| | | | | | Circle | 0.955 | 0.403 | 0.321 | 0.472 | 0.713 | 0.938 | 0.683 | 0.732 |
| | | | | | Star | 0.976 | 0.386 | 0.313 | 0.445 | 0.727 | 0.979 | 0.778 | 0.798 |
| | | | | | Dense | 0.030 | – | – | 0.055 | 0.699 | 0.978 | 0.755 | 0.773 |

Table S6: Full recovery results for the compositional outcome when varying $T$. Values are averaged over 100 simulation replicates. Boldface indicates the numerically largest average MCC across the structures for which MCC is defined.

| $N$ | $T$ | $p$ | $B$ | Rep. | Structure | Metrics for $\Theta$ | | | | Metrics for $\Gamma$ | | | |
|---|---|---|---|---|---|---|---|---|---|---|---|---|---|
| | | | | | | SEN | SPE | MCC | F1 | SEN | SPE | MCC | F1 |
| 100 | 10 | 10 | 100 | 100 | AR1 | 0.883 | 0.723 | 0.502 | 0.576 | 0.678 | 0.956 | 0.699 | 0.738 |
| | | | | | AR2 | 0.626 | 0.718 | 0.343 | 0.584 | 0.730 | 0.956 | 0.738 | 0.775 |
| | | | | | Block | 0.891 | 0.920 | **0.815** | 0.885 | 0.684 | 0.959 | 0.712 | 0.748 |
| | | | | | Circle | 0.911 | 0.402 | 0.282 | 0.454 | 0.633 | 0.856 | 0.478 | 0.582 |
| | | | | | Star | 0.857 | 0.486 | 0.289 | 0.438 | 0.504 | 0.981 | 0.617 | 0.630 |
| | | | | | Dense | 0.031 | – | – | 0.058 | 0.749 | 0.953 | **0.752** | 0.788 |
| 100 | 30 | 10 | 100 | 100 | AR1 | 0.880 | 0.730 | 0.508 | 0.587 | 0.567 | 0.974 | 0.650 | 0.674 |
| | | | | | AR2 | 0.636 | 0.764 | 0.402 | 0.610 | 0.758 | 0.956 | **0.766** | 0.801 |
| | | | | | Block | 0.888 | 0.920 | **0.813** | 0.882 | 0.619 | 0.974 | 0.693 | 0.718 |
| | | | | | Circle | 0.908 | 0.382 | 0.265 | 0.445 | 0.575 | 0.923 | 0.544 | 0.617 |
| | | | | | Star | 0.867 | 0.563 | 0.369 | 0.494 | 0.579 | 0.976 | 0.665 | 0.688 |
| | | | | | Dense | 0.062 | – | – | 0.113 | 0.576 | 0.981 | 0.677 | 0.692 |
| 100 | 50 | 10 | 100 | 100 | AR1 | 0.927 | 0.682 | 0.494 | 0.574 | 0.726 | 0.959 | 0.746 | 0.780 |
| | | | | | AR2 | 0.613 | 0.781 | 0.398 | 0.603 | 0.801 | 0.940 | 0.770 | 0.809 |
| | | | | | Block | 0.823 | 0.992 | **0.838** | 0.866 | 0.729 | 0.961 | 0.754 | 0.786 |
| | | | | | Circle | 0.918 | 0.405 | 0.291 | 0.458 | 0.779 | 0.709 | 0.423 | 0.549 |
| | | | | | Star | 0.899 | 0.692 | 0.511 | 0.597 | 0.765 | 0.945 | 0.748 | 0.788 |
| | | | | | Dense | 0.003 | – | – | 0.006 | 0.738 | 0.966 | **0.769** | 0.799 |

Table S7: Full recovery results for the binary outcome when varying $p$. Values are averaged over 100 simulation replicates. Boldface indicates the numerically largest average MCC across the structures for which MCC is defined.

| $N$ | $T$ | $p$ | $B$ | Rep. | Structure | Metrics for $\Theta$ | | | | Metrics for $\Gamma$ | | | |
|---|---|---|---|---|---|---|---|---|---|---|---|---|---|
| | | | | | | SEN | SPE | MCC | F1 | SEN | SPE | MCC | F1 |
| 100 | 10 | 10 | 100 | 100 | AR1 | 0.926 | 0.408 | 0.284 | 0.432 | 0.585 | 0.933 | 0.558 | 0.625 |
| | | | | | AR2 | 0.811 | 0.429 | 0.249 | 0.588 | 0.480 | 0.974 | 0.563 | 0.590 |
| | | | | | Block | 0.838 | 0.453 | **0.313** | 0.664 | 0.551 | 0.950 | 0.563 | 0.614 |
| | | | | | Circle | 0.869 | 0.408 | 0.245 | 0.441 | 0.542 | 0.889 | 0.443 | 0.546 |
| | | | | | Star | 0.844 | 0.376 | 0.190 | 0.389 | 0.476 | 0.969 | 0.547 | 0.576 |
| | | | | | Dense | 0.637 | – | – | 0.775 | 0.508 | 0.969 | **0.575** | 0.604 |
| 100 | 10 | 30 | 100 | 100 | AR1 | 0.889 | 0.871 | 0.498 | 0.487 | 0.603 | 0.880 | 0.469 | 0.578 |
| | | | | | AR2 | 0.589 | 0.886 | 0.424 | 0.504 | 0.732 | 0.828 | 0.499 | 0.606 |
| | | | | | Block | 0.251 | 0.895 | 0.193 | 0.367 | 0.469 | 0.919 | 0.426 | 0.522 |
| | | | | | Circle | 0.928 | 0.868 | **0.521** | 0.505 | 0.647 | 0.843 | 0.451 | 0.569 |
| | | | | | Star | 0.292 | 0.850 | 0.096 | 0.170 | 0.607 | 0.899 | 0.504 | 0.603 |
| | | | | | Dense | 0.129 | – | – | 0.228 | 0.640 | 0.910 | **0.550** | 0.639 |
| 100 | 10 | 50 | 100 | 100 | AR1 | 0.814 | 0.913 | 0.448 | 0.422 | 0.530 | 0.873 | 0.397 | 0.520 |
| | | | | | AR2 | 0.500 | 0.920 | 0.361 | 0.414 | 0.593 | 0.904 | 0.503 | 0.601 |
| | | | | | Block | 0.142 | 0.919 | 0.097 | 0.230 | 0.433 | 0.891 | 0.343 | 0.463 |
| | | | | | Circle | 0.860 | 0.908 | **0.466** | 0.433 | 0.521 | 0.883 | 0.405 | 0.523 |
| | | | | | Star | 0.163 | 0.907 | 0.046 | 0.095 | 0.542 | 0.895 | 0.444 | 0.552 |
| | | | | | Dense | 0.095 | – | – | 0.173 | 0.631 | 0.862 | 0.464 | 0.578 |

Table S8: Full recovery results for the count outcome when varying $p$. Values are averaged over 100 simulation replicates. Boldface indicates the numerically largest average MCC across the structures for which MCC is defined.

| $N$ | $T$ | $p$ | $B$ | Rep. | Structure | Metrics for $\Theta$ | | | | Metrics for $\Gamma$ | | | |
|---|---|---|---|---|---|---|---|---|---|---|---|---|---|
| | | | | | | SEN | SPE | MCC | F1 | SEN | SPE | MCC | F1 |
| 100 | 10 | 10 | 100 | 100 | AR1 | 0.954 | 0.636 | 0.478 | 0.562 | 0.718 | 0.926 | 0.665 | 0.716 |
| | | | | | AR2 | 0.802 | 0.703 | 0.494 | 0.698 | 0.766 | 0.963 | **0.776** | 0.805 |
| | | | | | Block | 0.943 | 0.932 | **0.876** | 0.929 | 0.663 | 0.956 | 0.684 | 0.716 |
| | | | | | Circle | 0.951 | 0.408 | 0.319 | 0.474 | 0.702 | 0.898 | 0.600 | 0.671 |
| | | | | | Star | 0.958 | 0.426 | 0.325 | 0.453 | 0.736 | 0.955 | 0.736 | 0.770 |
| | | | | | Dense | 0.101 | – | – | 0.179 | 0.714 | 0.960 | 0.732 | 0.761 |
| 100 | 10 | 30 | 100 | 100 | AR1 | 0.985 | 0.885 | 0.576 | 0.550 | 0.616 | 0.968 | 0.660 | 0.703 |
| | | | | | AR2 | 0.712 | 0.910 | 0.558 | 0.617 | 0.731 | 0.976 | **0.765** | 0.800 |
| | | | | | Block | 0.830 | 0.998 | **0.844** | 0.904 | 0.594 | 0.963 | 0.630 | 0.678 |
| | | | | | Circle | 0.993 | 0.865 | 0.551 | 0.522 | 0.665 | 0.966 | 0.689 | 0.735 |
| | | | | | Star | 0.992 | 0.948 | 0.745 | 0.739 | 0.657 | 0.966 | 0.687 | 0.731 |
| | | | | | Dense | 0.000 | – | – | 0.000 | 0.641 | 0.985 | 0.721 | 0.749 |
| 100 | 10 | 50 | 100 | 100 | AR1 | 0.853 | 0.977 | 0.707 | 0.708 | 0.580 | 0.971 | 0.635 | 0.681 |
| | | | | | AR2 | 0.610 | 0.959 | 0.550 | 0.586 | 0.698 | 0.976 | **0.739** | 0.778 |
| | | | | | Block | 0.705 | 0.993 | **0.732** | 0.823 | 0.518 | 0.970 | 0.586 | 0.630 |
| | | | | | Circle | 1.000 | 0.900 | 0.519 | 0.461 | 0.617 | 0.976 | 0.677 | 0.718 |
| | | | | | Star | 0.388 | 0.999 | 0.477 | 0.434 | 0.588 | 0.976 | 0.657 | 0.696 |
| | | | | | Dense | 0.005 | – | – | 0.009 | 0.588 | 0.989 | 0.692 | 0.717 |

Table S9: Full recovery results for the compositional outcome when varying $p$. Values are averaged over 100 simulation replicates. Boldface indicates the numerically largest average MCC across the structures for which MCC is defined.

| $N$ | $T$ | $p$ | $B$ | Rep. | Structure | Metrics for $\Theta$ | | | | Metrics for $\Gamma$ | | | |
|---|---|---|---|---|---|---|---|---|---|---|---|---|---|
| | | | | | | SEN | SPE | MCC | F1 | SEN | SPE | MCC | F1 |
| 100 | 10 | 10 | 100 | 100 | AR1 | 0.883 | 0.723 | 0.502 | 0.576 | 0.678 | 0.956 | 0.699 | 0.738 |
| | | | | | AR2 | 0.626 | 0.718 | 0.343 | 0.584 | 0.730 | 0.956 | 0.738 | 0.775 |
| | | | | | Block | 0.891 | 0.920 | **0.815** | 0.885 | 0.684 | 0.959 | 0.712 | 0.748 |
| | | | | | Circle | 0.911 | 0.402 | 0.282 | 0.454 | 0.633 | 0.856 | 0.478 | 0.582 |
| | | | | | Star | 0.857 | 0.486 | 0.289 | 0.438 | 0.504 | 0.981 | 0.617 | 0.630 |
| | | | | | Dense | 0.031 | – | – | 0.058 | 0.749 | 0.953 | **0.752** | 0.788 |
| 100 | 10 | 30 | 100 | 100 | AR1 | 0.765 | 0.930 | **0.543** | 0.539 | 0.431 | 0.980 | 0.548 | 0.568 |
| | | | | | AR2 | 0.429 | 0.936 | 0.383 | 0.437 | 0.618 | 0.958 | 0.641 | 0.694 |
| | | | | | Block | 0.424 | 0.874 | 0.334 | 0.532 | 0.480 | 0.965 | 0.547 | 0.590 |
| | | | | | Circle | 0.944 | 0.835 | 0.479 | 0.450 | 0.573 | 0.944 | 0.576 | 0.640 |
| | | | | | Star | 0.442 | 0.957 | 0.398 | 0.423 | 0.444 | 0.974 | 0.538 | 0.569 |
| | | | | | Dense | 0.013 | – | – | 0.024 | 0.533 | 0.988 | **0.652** | 0.671 |
| 100 | 10 | 50 | 100 | 100 | AR1 | 0.837 | 0.936 | **0.522** | 0.496 | 0.390 | 0.979 | 0.508 | 0.528 |
| | | | | | AR2 | 0.221 | 0.980 | 0.274 | 0.279 | 0.479 | 0.964 | 0.540 | 0.591 |
| | | | | | Block | 0.173 | 0.964 | 0.194 | 0.265 | 0.352 | 0.966 | 0.433 | 0.472 |
| | | | | | Circle | 0.884 | 0.907 | 0.474 | 0.431 | 0.446 | 0.956 | 0.491 | 0.550 |
| | | | | | Star | 0.141 | 0.991 | 0.222 | 0.202 | 0.384 | 0.980 | 0.504 | 0.522 |
| | | | | | Dense | 0.000 | – | – | 0.000 | 0.410 | 0.988 | **0.555** | 0.561 |

### S5. Convergence diagnostics

In this section, we aim to assess whether the estimates obtained under the tuning parameters selected in the pilot stage stabilize over the Monte Carlo EM iterations. For each combination of outcome type and contemporaneous dependence structure, a tuning parameter pair $(\lambda_\Gamma, \lambda_\Theta)$ was selected using a pilot run with a fixed random seed. We then examined the iteration histories of the corresponding estimates $\widehat{\Gamma}^{(k)}$ and $\widehat{\Theta}^{(k)}$. After confirming that the estimates entered a stable numerical regime, the selected tuning parameter pair was held fixed and applied to the 100 independent simulation replicates used to compute the support-recovery results.

At iteration $k$, the stability of the estimates is measured using the relative Frobenius changes

$$\Delta_\Gamma^{(k)} = \frac{\left\|\widehat{\Gamma}^{(k)} - \widehat{\Gamma}^{(k-1)}\right\|_\mathrm{F}}{\left\|\widehat{\Gamma}^{(k-1)}\right\|_\mathrm{F} + 10^{-10}}, \qquad \Delta_\Theta^{(k)} = \frac{\left\|\widehat{\Theta}^{(k)} - \widehat{\Theta}^{(k-1)}\right\|_\mathrm{F}}{\left\|\widehat{\Theta}^{(k-1)}\right\|_\mathrm{F} + 10^{-10}}.$$

Smaller values indicate that successive estimates are changing less and have entered a more stable

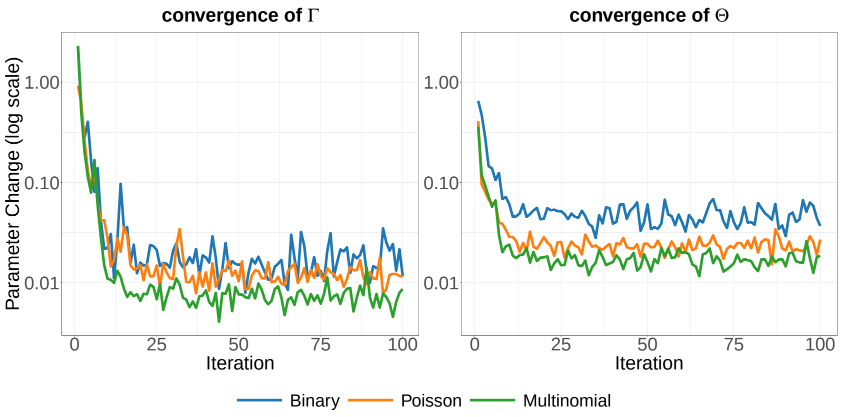

(a) AR1

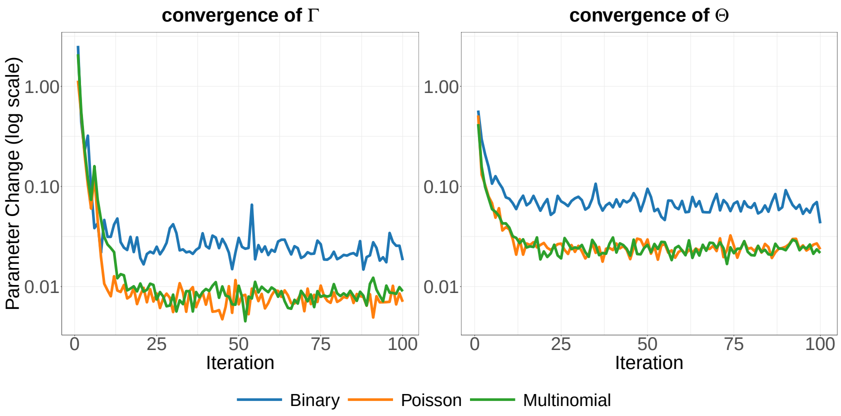

(b) AR2

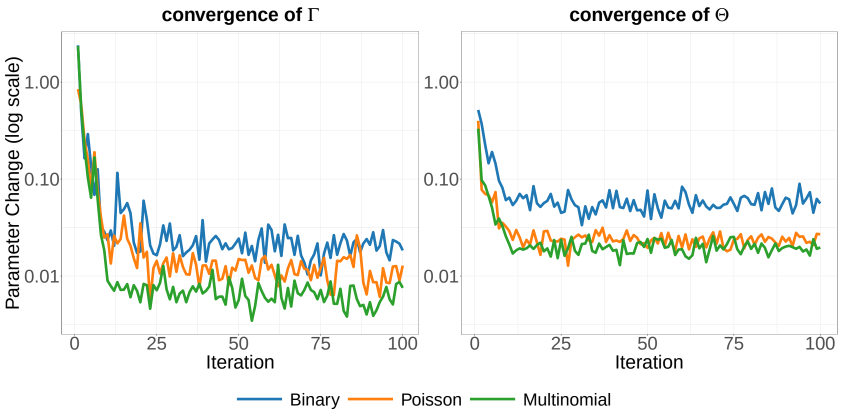

(c) Block

Fig. S2: Convergence diagnostics for the pilot tuning runs under the baseline setting: AR1, AR2, and Block structures.

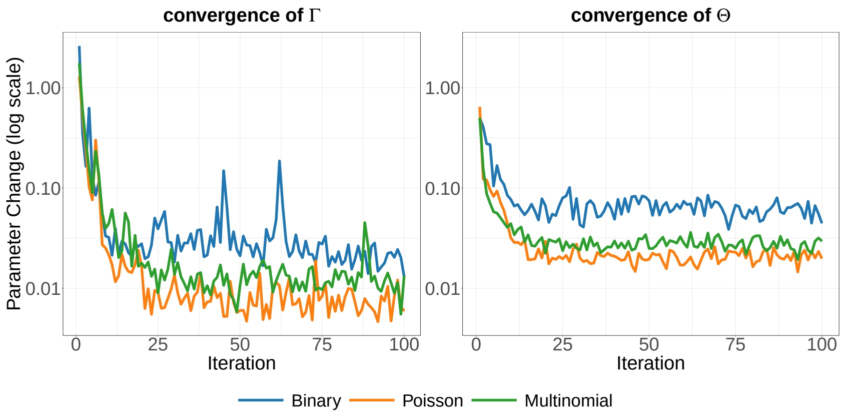

(a) Circle

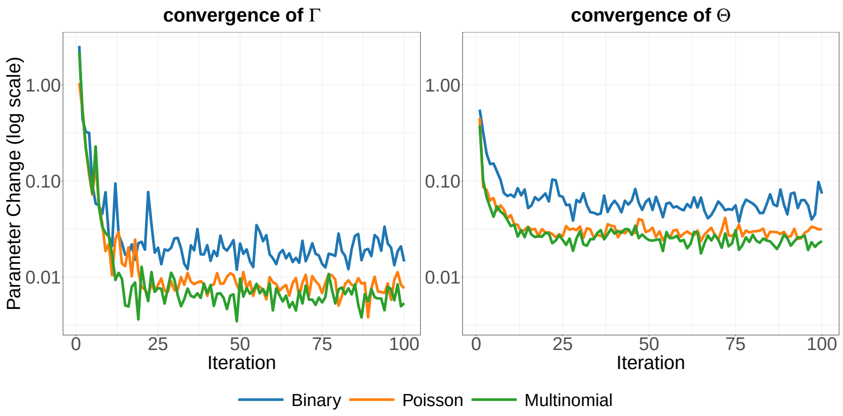

(b) Star

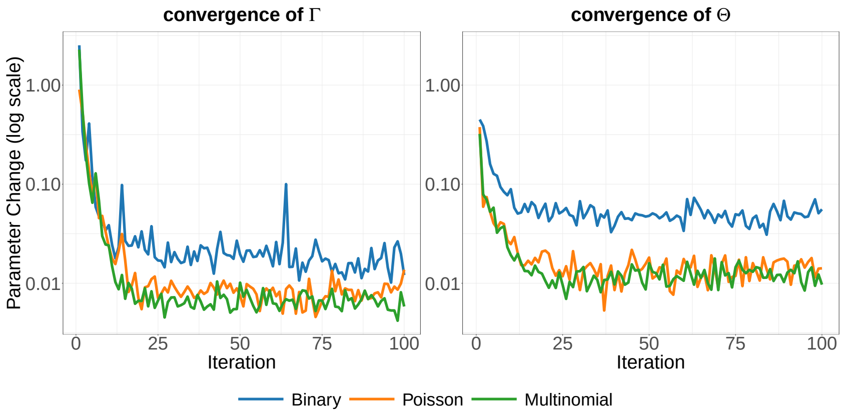

(c) Dense

Fig. S3: Convergence diagnostics for the pilot tuning runs under the baseline setting: Circle, Star, and Dense structures.

numerical regime.

Figures S2 and S3 show these relative changes under the baseline setting $(N, T, p, B) = (100, 10, 10, 100)$. Results are shown separately for the six contemporaneous dependence structures and for the binary, count, and compositional outcomes. The left panel for each structure displays $\Delta_\Gamma^{(k)}$, and the right panel displays $\Delta_\Theta^{(k)}$. The vertical axes are shown on a logarithmic scale. Across all six structures, the changes in both $\widehat{\Gamma}$ and $\widehat{\Theta}$ decrease sharply during the initial iterations and subsequently fluctuate within a much smaller and relatively stable range. The remaining nonzero fluctuations are expected because the Monte Carlo E-step introduces iteration-to-iteration variability.

The binary fits generally show larger fluctuations than the count and compositional fits, particularly for $\widehat{\Theta}$. Nevertheless, the binary estimates also enter a stable range after the initial iterations. Overall, the diagnostics indicate that the estimates associated with the selected pilot tuning parameters exhibit stable numerical behavior across the six dependence structures. The selected tuning parameter pairs were therefore fixed and used in the subsequent simulation replicates.
